# A groove brain-music interface for enhancing individual experience of urge to move

**DOI:** 10.1101/2025.09.16.676544

**Authors:** Takahide Etani, Sotaro Kondoh, Yuna Sakakibara, Keigo Yoshida, Yasushi Naruse, Yasuhiko Imamura, Takuya Ibaraki, Shinya Fujii

## Abstract

When we listen to music, we often feel a pleasurable urge to move to music, known as groove. While previous studies have identified musical features that contribute to the groove experience, such as syncopation and tempo, they also report individual differences in which kinds of music people experience groove. Therefore, recommending groove-eliciting music requires accounting for individual differences. In this study, we aimed to develop a groove brain-music interface (G-BMI) that generates personalized playlists to maximize each individual’s groove experience, using a neurofeedback system based on in-ear EEG. Twenty-four participants listened to three high-groove and three low-groove musical excerpts and rated their “urge to move.” Using these ratings and the recorded EEG, we trained two LASSO models to build the G-BMI. Model 1 predicted urge to move from acoustic features extracted with VGGish, a pretrained neural network. Model 2 classified EEG data as recorded during listening to high-groove or low-groove music. Using Model 1, we ranked 7,225 candidate songs by predicted groove and assembled one groove-augmenting and one groove-diminishing playlist. Using Models 1 and 2, we created two additional playlists that updated Model 1 and the ranking in real-time based on in-ear EEG. Participants then listened to all four playlists and rated them on items including “urge to move.” The groove-augmenting playlist that incorporated EEG achieved the highest “urge to move” ratings. These findings suggest that a personalized neurofeedback system employing EEG can help maximize individual groove experience.

## 1. Introduction

### 1.1 Groove

When we listen to music, we often feel a pleasurable urge to move in time with it, which is known as groove (Etani et al. 2024; Janata et al. 2012; Witek et al. 2014; Senn et al. 2023; Madison 2006; Zalta et al. 2024), more recently termed PLUMM (the *pleasurable urge to move to music*) (Matthews et al., 2023). Which kinds of music most strongly elicit the groove experience has long been debated. For example, ethnomusicologist Charles Keil argued that participatory discrepancies, subtle temporal deviations such as asynchronies between instruments and co-performers that are not captured in standard notation, are important for groove (Keil & Feld, 1994).

In recent years, research on the psychology and neuroscience of groove has grown rapidly. Complementing the deep scrutiny by scholars, these have revealed musical features related to the groove experience, including the degree of syncopation, beat salience, event density, tempo, bass, and harmony (Etani et al., 2018; Jerjen et al., 2024; Madison et al., 2011; Matthews et al., 2019; Senn et al., 2018; Witek et al., 2014). Beyond identifying musical features important for groove (Etani et al., 2024), there have also been attempts to generate groove-inducing audio. For example, Kawai et al. (2024) reproduced the groove of Jeff Porcaro’s hi-hat drumming using an oscillation driven reservoir computing model, a type of recurrent neural network in which firing rate units receive multiple sinusoidal oscillator inputs. Reservoir computing supports temporal information processing through neural dynamics and has been applied to modeling the dynamics of the cerebellum and basal ganglia. In that study, the authors trained the reservoir on Porcaro’s hi-hat performance and generated hi-hat patterns by adjusting event timing and amplitude to match his playing. Because such models emulate neural oscillatory dynamics, incorporating brain activity measures, such as EEG, may further aid groove generation.

While there are common features that generally induce the groove experience, there are also individual factors such as musical and dance sophistication modulate the extent to which people experience groove (Cameron et al. 2022; O’Connell et al. 2022; Senn et al. 2018; Stupacher et al. 2025; Pando-Naude et al. 2024; Benson et al. 2024). O’Connell et al. (2022) investigated whether music and dance sophistication influence the groove experience using the Goldsmiths Musical Sophistication Index (Gold-MSI) (Müllensiefen et al., 2014) and the Goldsmiths Dance Sophistication Index (Gold-DSI) (Rose et al., 2022). They found that scores on the Perceptual Abilities and Musical Training subscales of the Gold-MSI, as well as the Social Dancing subscale of the Gold-DSI, significantly affected the groove experience. In another study, Matthews et al. (2022) reported no significant difference in groove ratings between musicians and non-musicians, but found that the inverted U-shaped relationship between groove and rhythmic complexity was more pronounced in musicians (Matthews et al., 2022). In other words, compared with non-musicians, musicians preferred rhythms with medium syncopation over those with low or high syncopation. A similar pattern was observed for dancers compared with non-dancers (Cameron et al., 2022). In addition, factors such as age, preferred musical style, and emotional responsiveness also influence the groove experience (Engel et al., 2022; Senn et al., 2018), indicating individual differences in both the types of music that elicit groove. Therefore, to maximize each listener’s groove experience, it is crucial to account for individual differences in addition to identifying common factors. Furthermore, because musical and dance training alter brain structure and function (e.g., Hyde et al. 2009; Schlaug et al. 2005; Karpati et al. 2015), using neural measures such as EEG may help maximize each individual’s groove experience.

### 1.2 Closed-loop feedback system

In recent years, particularly alongside advances in AI, interest has grown in personalized services and related research, for example in precision and personalized medicine (Delpierre & Lefèvre, 2023). One effective approach in this context is the use of closed-loop feedback systems. In closed-loop feedback, information about the individual (such as hormonal and physiological data) is obtained in real-time, and the output is continuously updated based on that information. A representative example is the insulin pump, which measures a patient’s blood glucose levels in real-time and adjusts the amount of insulin delivered accordingly (Berget et al., 2019). More recently, the development of closed-loop feedback systems that utilize neural activity has advanced as well. Scangos et al. (2021) identified biomarkers related to depressive states in patients with treatment resistant depression, and based on that information, developed a closed-loop deep brain stimulation system that delivered stimulation accordingly and alleviated symptoms (Scangos et al., 2021).

### 1.3 Music and closed-loop feedback system

In the domain of music, there have also been attempts to use closed-loop systems to maximize individual musical experience. Trost et al. (2024) developed a system that increases activity in the amygdala by delivering fMRI feedback to the performer in real-time (Trost et al., 2024). In this study, the listener remained in the MRI scanner and listened either to a recording of a piano performance (recording condition) or to a live performance by a pianist who received real-time feedback about the listener’s amygdala activity (live condition). The authors found that amygdala activity was stronger in the live condition than in the recording condition, suggesting that the neurofeedback system can enhance the listener’s musical experience. Other recent studies have developed closed-loop neurofeedback systems that use each participant’s EEG to maximize subjective responses during music listening, including nostalgia, well-being, and memory vividness (Sakakibara et al. 2025) as well as chills and pleasure (Kondoh et al., 2024). In another study, Awad et al. (2024) developed a device that uses feedback to deliver individualized music and thereby maximize gait assistance (Awad et al., 2024), which was approved by the FDA as a therapeutic tool for post stroke patients in 2025. This study indicates that it may also be possible to develop a feedback system that maximizes individual groove experience, enhancing the urge to move. Although closed-loop neurofeedback systems that maximize individual musical experiences have been developed recently, no system has yet been designed to maximize the groove experience.

### 1.4 Aims

Therefore, in this study, we aimed to develop a groove brain–music interface (G-BMI) that generates personalized playlists to maximize each individual’s groove experience, based on in-ear EEG data.

## 2. Methods

### 2.1 Overview of the study

To build the G-BMI, we followed three steps: data recording, modeling, and playlist generation (see Supplementary Material for details).

#### 2.1.1 Recording

To obtain EEG data during music listening, we drew on a previous stimulus set (Janata et al., 2012). From a pool of seven high-groove and seven low-groove excerpts, we selected three high-groove and three low-groove excerpts (90 seconds each) for each participant (Table 1). After listening to each musical excerpt, participants rated their “urge to move” and “pleasure” on a visual analog scale (VAS) ranging from 0 to 100. In-ear EEG was recorded throughout listening using an in-ear EEG device (VIE Inc., Japan).

**Table 1:** Musical excerpts used for the experiment selected from Janata et al. (2012).

| High-groove music |  |  |
| --- | --- | --- |
| Artist | Title | Groove rating (ranking) |
| Stevie Wonder | Superstition | 108.7 (#1) |
| FH1 (Funky Hobo #1) | Its a wrap (Bye, bye) | 105.9 (#2) |

| Parliament | Flash light | 105.1 (#3) |
| --- | --- | --- |
| LaBelle | Lady Marmalade | 102.5 (#4) |
| The Clinton Administration | Up for the downstroke | 102.4 (#5) |
| Funk Squad | Mama Cita | 101.6 (#6) |
| Leela James | Music | 101.1 (#7) |
| Low-groove music |  |  |
| Artist | Title | Groove rating |
| William Coulter and Friends | Citi Na GCumman | 25.2 (#142) |
| Dean Magraw | Dawn star | 34.8 (#143) |
| Kaki King | Fortuna | 32.6 (#144) |
| The Gabe Dixon Band | Beauty of the sea | 32.1 (#145) |
| Alison Brown | Sweet thing | 30.9 (#146) |
| Barry Phillips | Thugamar Fin an Samhradh Linn | 29.3 (#148) |
| Adrian Legg | Hymn for Jaco | 29.3 (#148) |

#### 2.1.2 Modeling

We developed two models. First, we extracted acoustic features of high-groove music using VGGish, a pretrained neural network, and fit a LASSO regression model to predict subjective “urge to move” ratings from these features (Model 1). Second, we trained a generalized LASSO classifier to distinguish EEG states associated with listening to high-groove versus low-groove music (Model 2).

#### 2.1.3 Generating playlists

Using Model 1 and Model 2, we created four playlists: AugEEG, AugNoEEG, DimNoEEG, and DimEEG. We first analyzed the acoustic features of 7,225 music excerpts using VGGish and ranked them by the predicted “urge to move” from Model 1. Tracks for the AugEEG and AugNoEEG playlists were selected randomly from the top songs of the ranking, whereas tracks for the DimEEG and DimNoEEG playlists were selected from the bottom. Each playlist contained seven excerpts (60 seconds each). The first track in each playlist was chosen from those ranked between 3,577 and 3,649 to ensure comparable starting points.

For the AugEEG and DimEEG playlists, in-ear EEG was recorded while participants listened. After each excerpt, Model 1 and the ranking were updated using the EEG-based “urge to move” predictions from Model 2 together with acoustic-feature predictions from VGGish. In contrast, neither Model 1 nor the ranking was updated for the AugNoEEG and DimNoEEG playlists.

Each participant listened to all four playlists in random order and rated each playlist on items related to groove, wellbeing, and emotion, including “urge to move” and “pleasure” (see Table 2 for the full list of rating items), using a visual analog scale (VAS) ranging from 0 to 100.

**Table 2:**
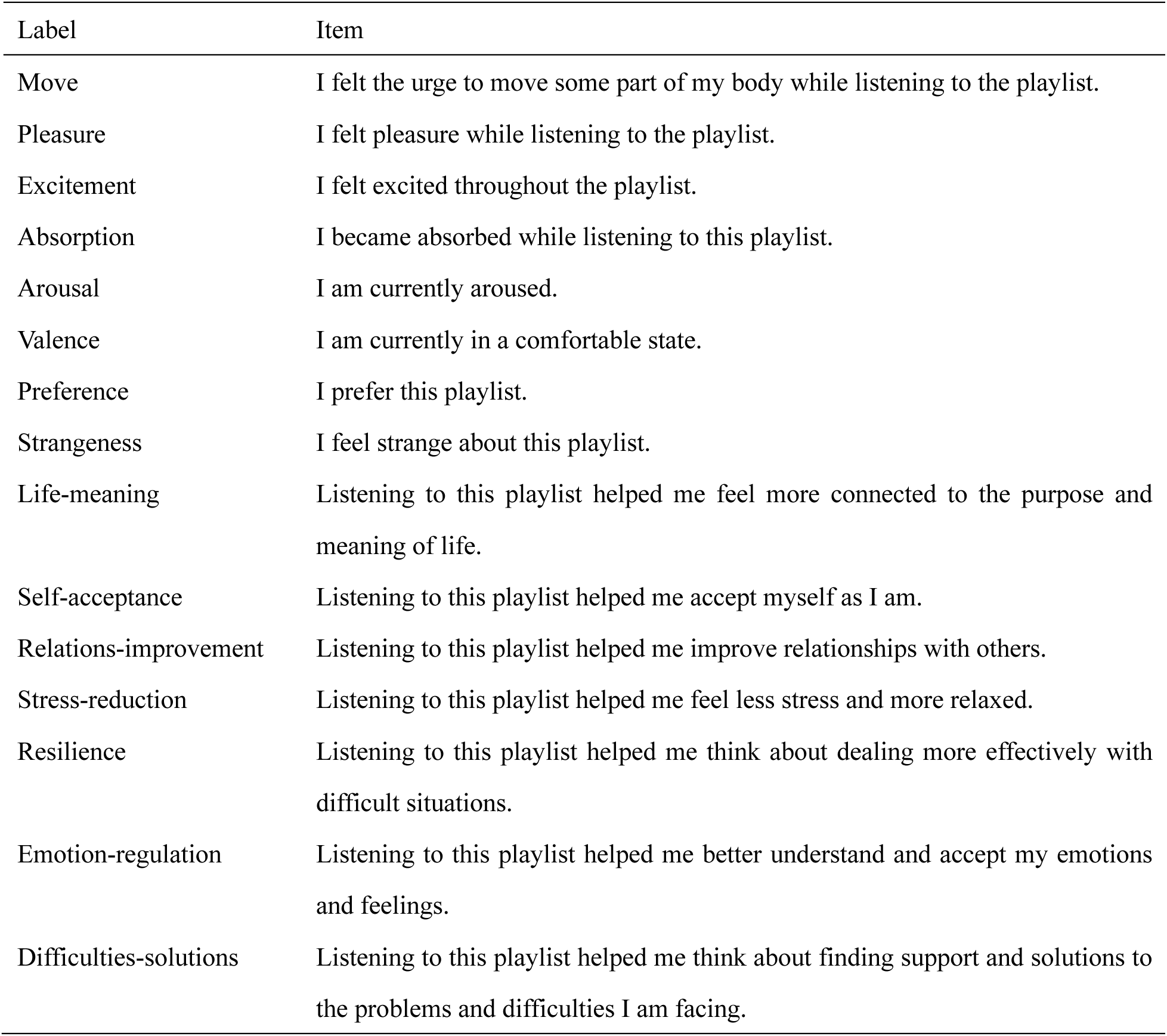
Rating items used for the experiment.

### 2.2 Participants

Twenty-four healthy participants took part in the study (mean age = 24.55 ± 5.91 years; 14 female participants). All participants received a verbal explanation and provided written informed consent. The study was approved by the Research Ethics Committee of Keio University Shonan Campus, Japan (No. 524). The sample size (*n* = 24) was determined a priori using G*Power 3 (Faul et al. 2007) for a one-way repeated measures analysis of variance (ANOVA), assuming a medium effect size, α = 0.05, and power = 0.8.

### 2.3 Procedure

Participants completed the recording session, during which they listened to three high-groove and three low-groove excerpts and rated each excerpt on “urge to move” and “pleasure” using a VAS (0 to 100). After model construction as described above, they listened to four playlists in random order and rated each playlist on 17 items (Table 2). Finally, they completed two questionnaires: the Gold-MSI and the Japanese version of the Barcelona Music Reward Questionnaire (BMRQ) (Sadakata et al. 2022; Müllensiefen et al. 2014). The entire experiment lasted approximately 90 minutes.

### 2.4 Statistical analyses

All statistical analyses were conducted using R version 3.6.3 and SPSS Statistics version 29.0.2.0.

#### 2.4.1 Subjective ratings of the music used for the training

We calculated mean VAS scores for “urge to move” and “pleasure” across the three high-groove and three low-groove excerpts. Normality was assessed with the Shapiro-Wilk test. Depending on the result, we compared the mean VAS scores of high-groove and low-groove music using either Student’s t test or the Wilcoxon signed rank test. The significance level was set at *p* = 0.05.

#### 2.4.2 Subjective ratings of the playlists

We excluded two participants from the VAS analyses because their data were not recorded due to a technical problem (*n* = 22). For the items “Life-meaning,” “Self-acceptance,” “Relations-improvement,” “Stress-reduction,” “Difficulty-solutions,” and “Emotion-regulation,” we excluded one additional participant due to missing responses (*n* = 21). For the decoded EEG analysis, we excluded four participants due to technical problems: EEG data were not recorded for one participant, and the EEG analysis failed for three participants (*n* = 20).

We calculated VAS scores of 17 items for each playlist. Normality was assessed with the Shapiro-Wilk test. For ratings that met normality, we conducted a one-way repeated measures ANOVA and, when the main effect was significant, followed up with pairwise comparisons. For ratings that violated normality for at least one playlist, we used the Friedman test and then paired Wilcoxon signed rank tests for pairwise comparisons. P-values for multiple testing were corrected using the Bonferroni method. As effect sizes, we reported partial eta-squared for the one-way ANOVA and Kendall’s W for the Friedman tests.

#### 2.4.3 Decoded urge to move

We fit a Bayesian generalized linear mixed model (GLMM) using the brms package in R to test whether decoded “urge to move” differed across the four playlists. The dependent variable was the baseline corrected “urge to move” decoded from in-ear EEG (“decoded urge to move”). Fixed effects were playlist (AugEEG, AugNoEEG, DimNoEEG, DimEEG) and within playlist song order (1 to 7). A random intercept was included for participant (Participant ID). The baseline was the first song. The GLMM was specified using the following formula:

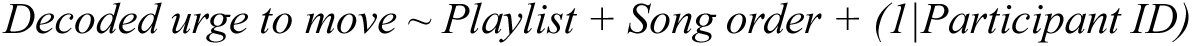

DimEEG was set as the reference level for all comparisons. We used the default noninformative priors in brms. No priors were set for the intercepts or standard deviations. Residual diagnostics indicated departures from normality, so we specified a Student’s t distribution for the residuals. Markov chain Monte Carlo sampling used four chains with 2,000 iterations per chain, including 1,000 warmup iterations, yielding 4,000 post warmup draws. Convergence was assessed with the potential scale reduction factor (R-hat). After fitting the GLMM, we conducted pairwise comparisons between playlists using the “emmeans” package (version 1.10.3). We obtained estimated marginal means and performed pairwise contrasts with 95% highest posterior density (HPD) intervals, and evaluated effects by whether the 95% HPD interval excluded zero.

#### 2.4.4 Individual differences (BMRQ and Gold-MSI)

To examine how individual differences affected the model’s ability to enhance “urge to move” ratings, we conducted a multiple regression analysis using subscales of the BMRQ and Gold-MSI. For each participant, we computed MoveDiffEEG, defined as the difference in “urge to move” ratings between AugEEG and DimEEG (AugEEG − DimEEG). We then performed stepwise multiple regression with MoveDiffEEG as the dependent variable and the following as independent variables: BMRQ factors (Emotion Evocation, Music Seeking, Sensory-Motor, Social Reward, Mood Regulation), the BMRQ total score, Gold-MSI subscales (Active Engagement, Perceptual Abilities, Musical Training, Singing Abilities, Emotions), and the Gold-MSI General Sophistication score. Model selection was based on the Akaike Information Criterion (AIC).

## 3. Results

### 3.1 Subjective ratings of the music used for the training

A t test on “urge to move” ratings showed that high-groove music received higher ratings than low-groove music (*t*_(21)_ = 8.92, *p* < 0.001, *d* = 0.89) (Figure 2). Likewise, a t test on “pleasure” ratings showed that high groove music received higher ratings than low groove music (*t*_(21)_ = 2.54, *p* = 0.019, *d* = 0.49) (Figure 3). These results confirmed our stimulus selection for model training.

**Figure 1:**
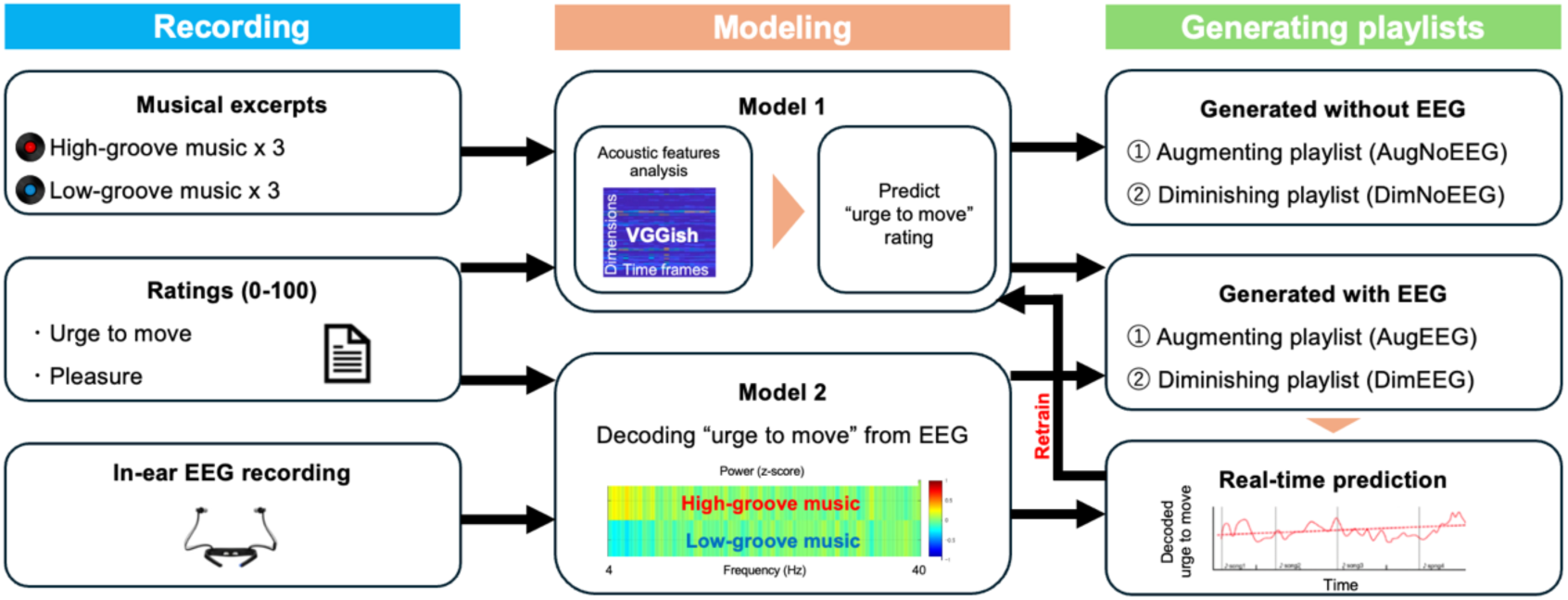
Overview of the study. From a pool of seven high-groove and seven low-groove excerpts, we selected three high-groove and three low-groove excerpts (90 seconds each) for each participant. After listening to each musical excerpt, participants rated their “urge to move” and “pleasure” on a visual analog scale (VAS) ranging from 0 to 100. In-ear EEG was recorded throughout listening using an in-ear EEG device. We developed two models. First, we extracted acoustic features of high-groove music using VGGish, a pretrained neural network, and fit a LASSO regression model to predict subjective “urge to move” ratings from these features (Model 1). Second, we trained a generalized LASSO classifier to distinguish EEG states associated with listening to high-groove versus low-groove music (Model 2). Using Model 1 and Model 2, we created four playlists: AugEEG, AugNoEEG, DimNoEEG, and DimEEG. We first analyzed the acoustic features of 7,225 music excerpts using VGGish and ranked them by the predicted “urge to move” from Model 1. Tracks for the AugEEG and AugNoEEG playlists were selected from the top of the ranking, whereas tracks for the DimEEG and DimNoEEG playlists were selected from the bottom. Each playlist contained seven excerpts (60 seconds each). For the AugEEG and DimEEG playlists, in-ear EEG was recorded while participants listened. After each excerpt, Model 1 and the ranking were updated using the EEG-based “urge to move” predictions from Model 2 together with acoustic-feature predictions from VGGish. In contrast, neither Model 1 nor the ranking was updated for the AugNoEEG and DimNoEEG playlists.

**Figure 2:**
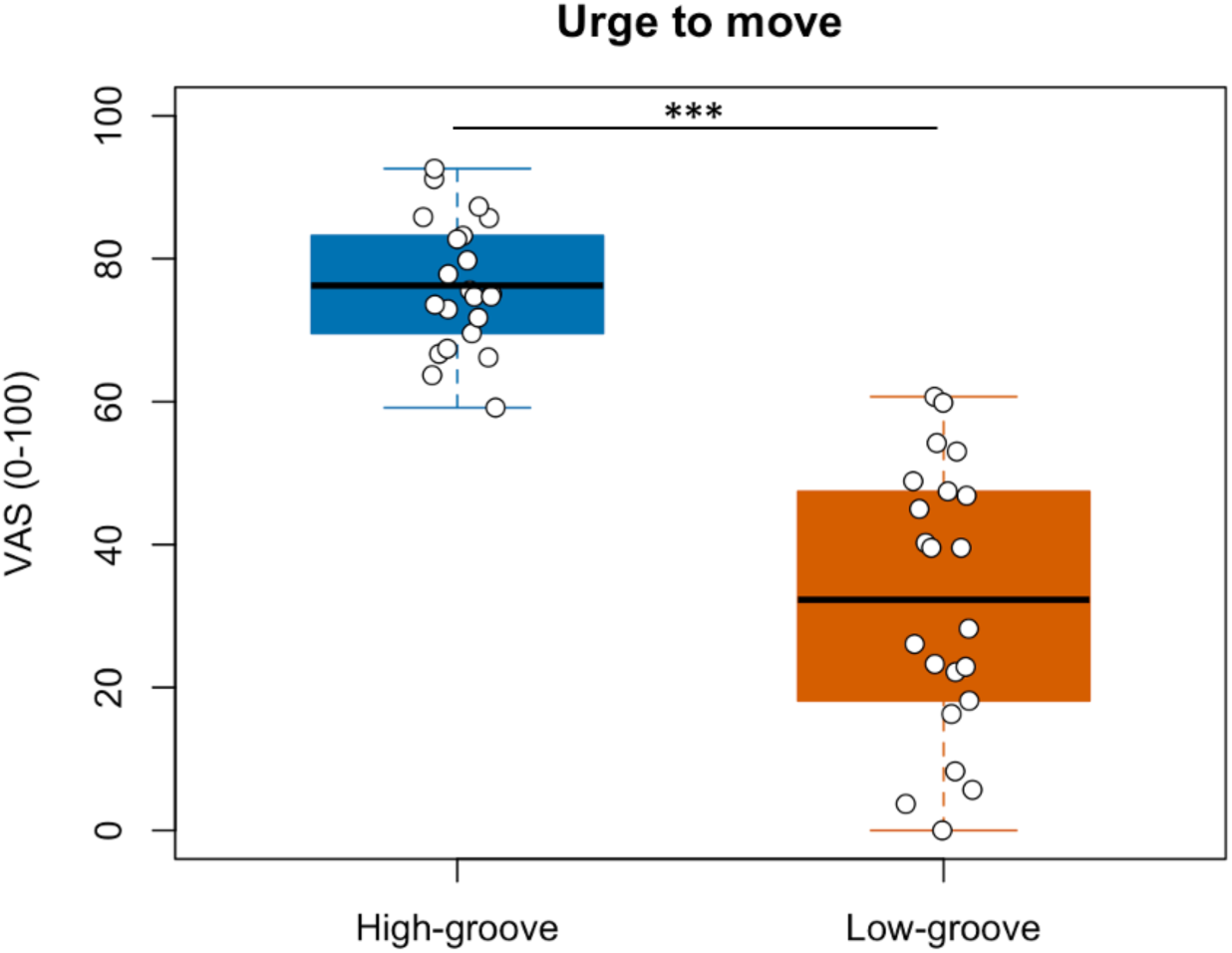
The results of the t test on the “urge to move” ratings. *** *p* < 0.001

**Figure 3:**
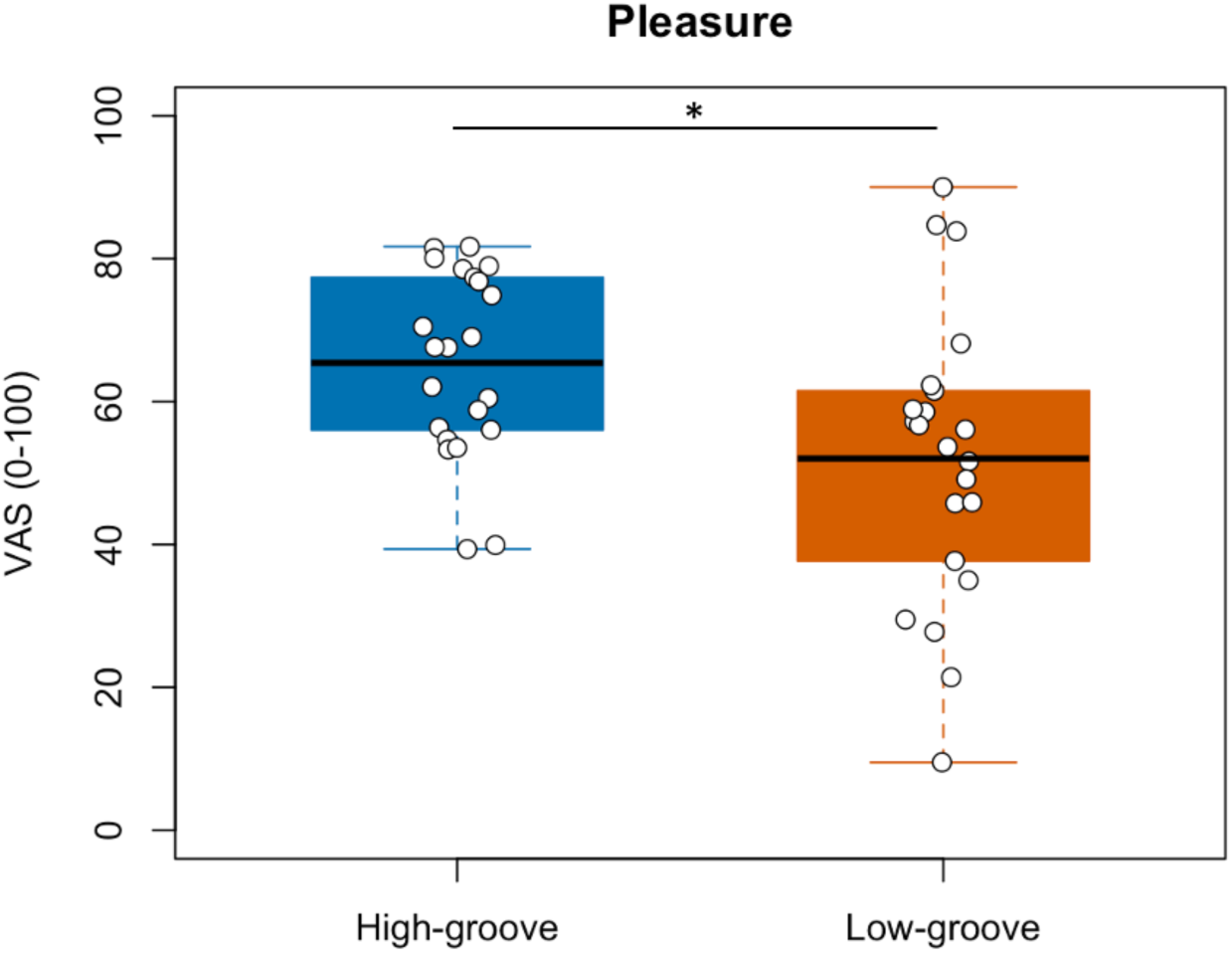
The results of the t test on the “pleasure” ratings. * *p* < 0.05

### 3.2 The accuracy of Model 2

The mean accuracy rates for the training and test sets were 81.1% and 68.6%, respectively, and the mean AUC for the trained and test data were 87.3% and 72.6%, respectively (see supplementary material for detail).

### 3.3 Subjective groove-related ratings for the playlists

The Friedman test revealed a significant difference in “urge to move” ratings across the four conditions (*χ²*(3) = 22.45, *p* < 0.001, with Kendall’s W = 0.34), indicating a moderate effect. Post hoc Wilcoxon signed rank tests with Bonferroni correction showed that ratings for AugEEG were significantly higher than those for DimNoEEG (*p* = 0.010) and DimEEG (*p* < 0.001) (Figure 4).

**Figure 4:**
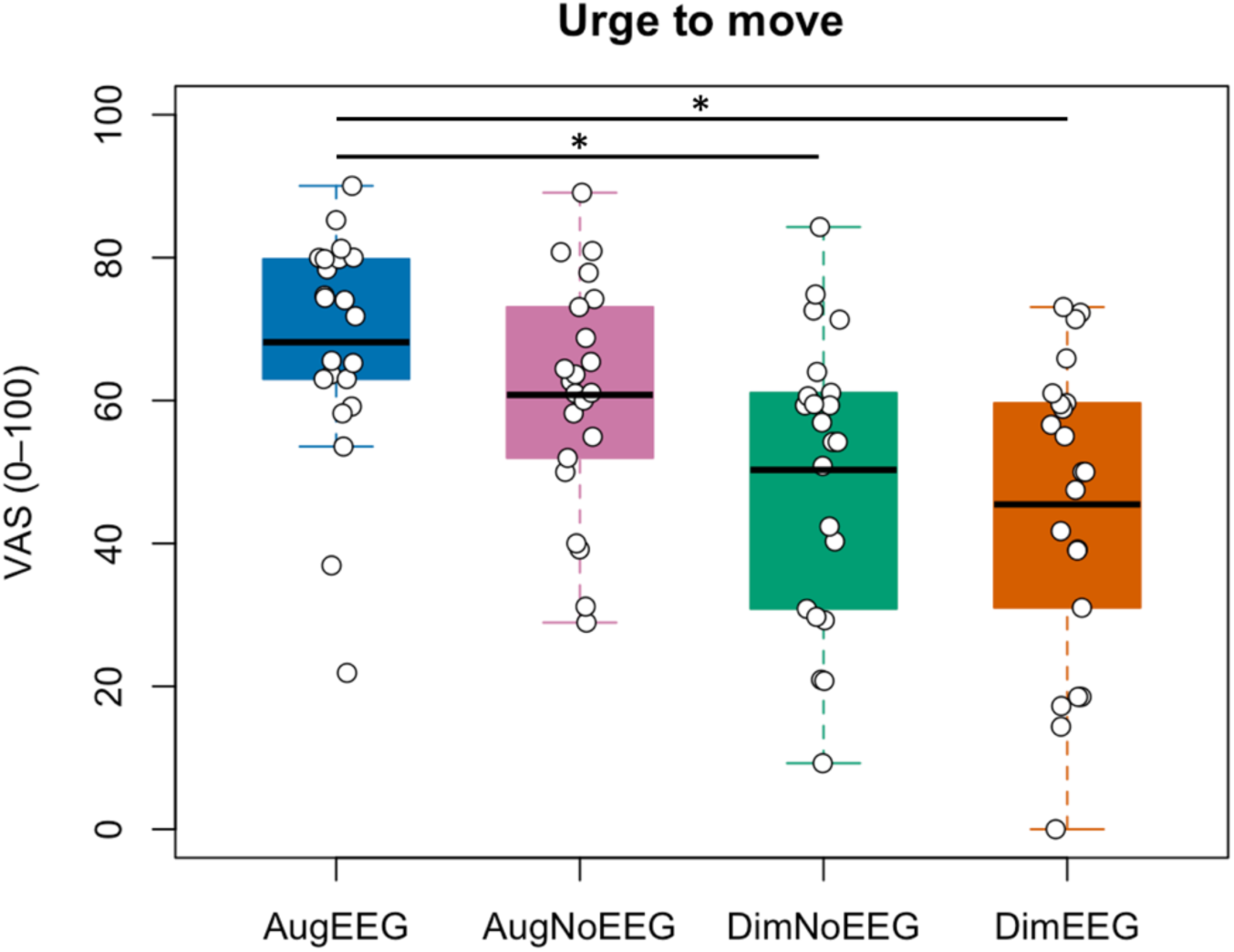
The results of the Friedman test on the “urge to move” ratings. * *p* < 0.05

The Friedman test showed no significant differences in “pleasure” ratings across the four conditions (*χ²*(3) = 3.00, *p* = 0.39, with Kendall’s W = 0.045) (Figure 5).

**Figure 5:**
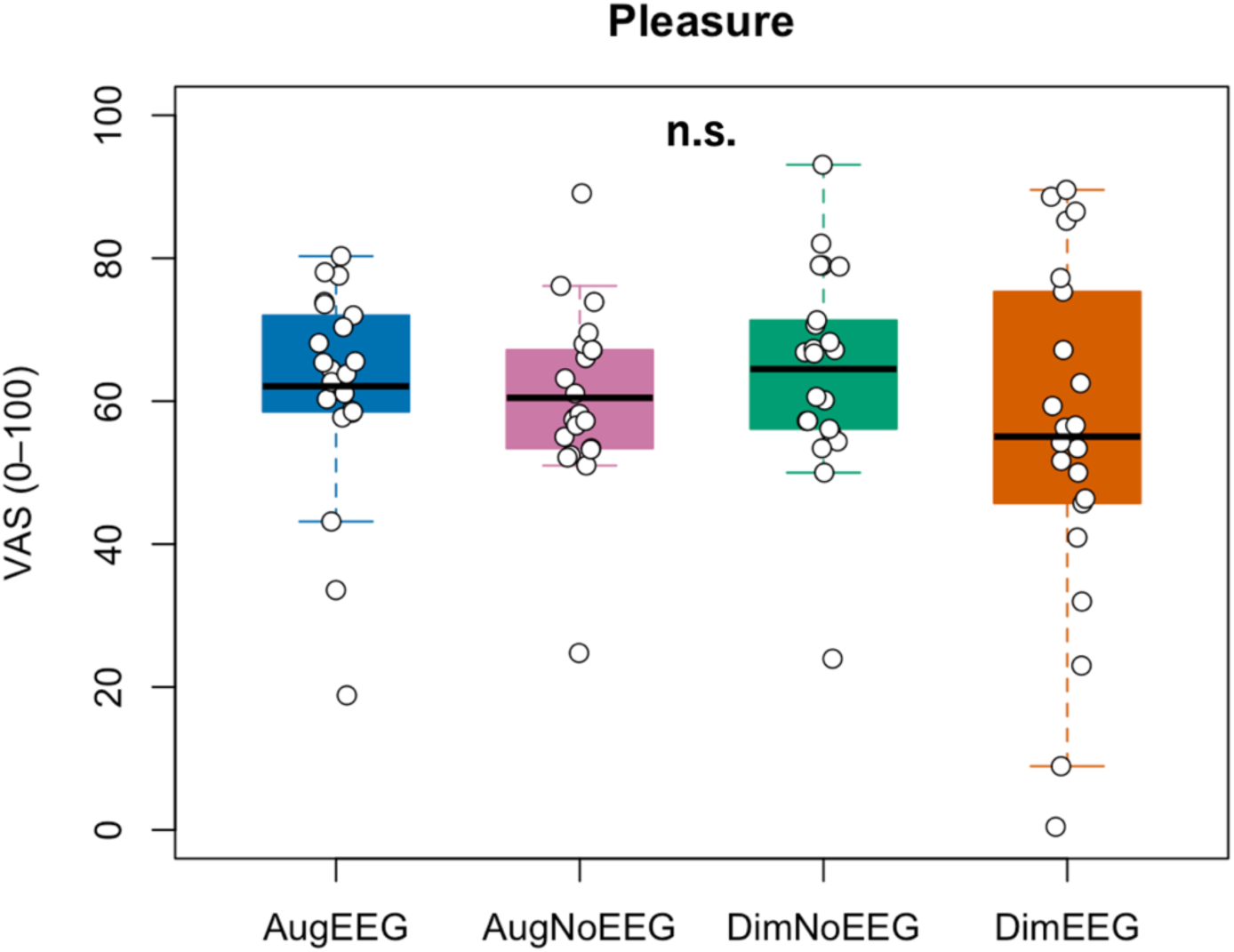
The results of the Friedman test on the “pleasure” ratings.

The Friedman test revealed a significant difference in “groove (urge to move + pleasure)” ratings across the four conditions (*χ²*(3) = 10.86, *p* = 0.013, with Kendall’s W = 0.16), indicating a small effect. Post hoc Wilcoxon signed rank tests with Bonferroni correction showed that ratings for AugEEG were significantly higher than those for DimEEG (*p* = 0.017) (Figure 6).

**Figure 6:**
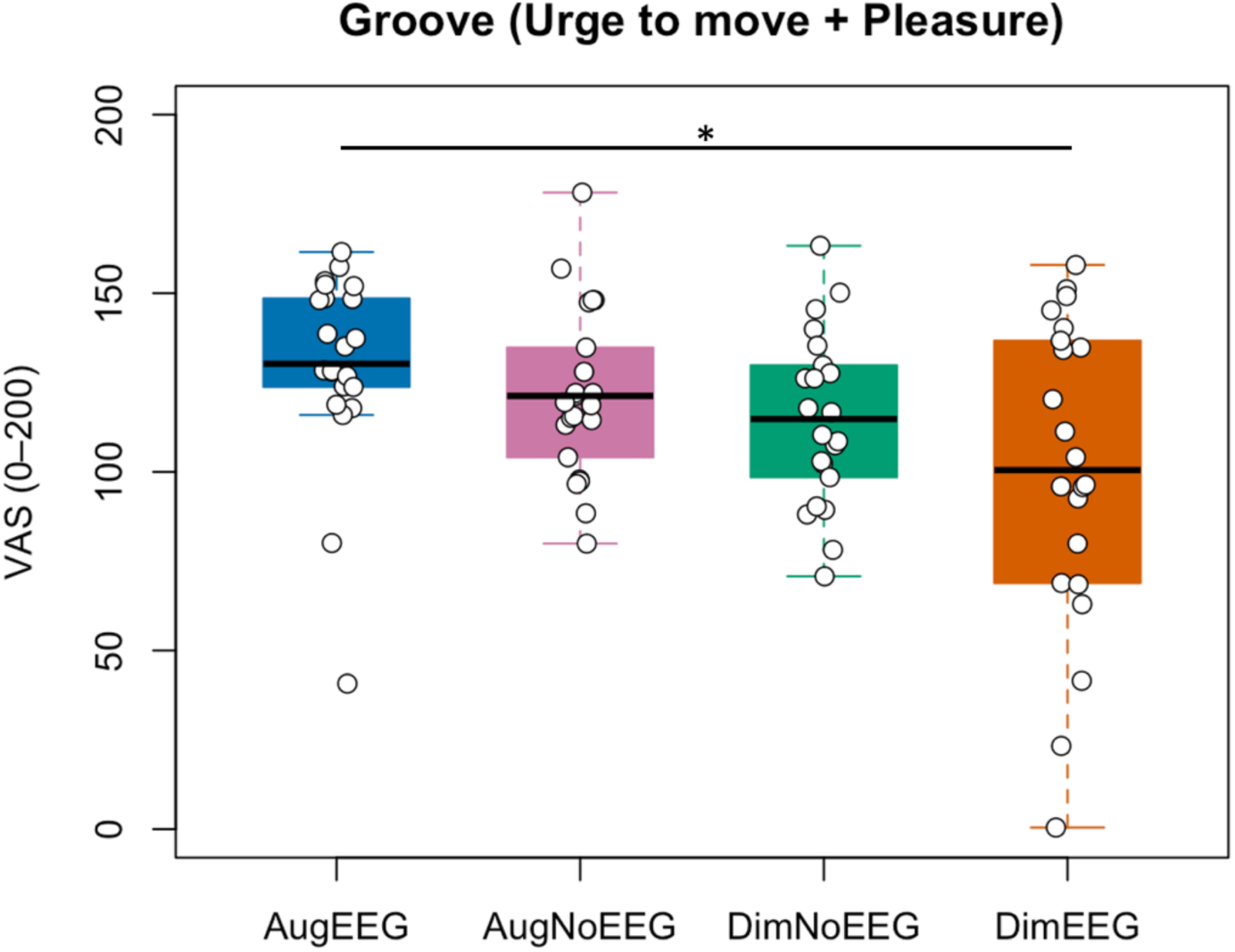
The results of the Friedman test on the “groove” ratings. * *p* < 0.05

### 3.4 Subjective well-being and emotion-related ratings for the playlists

We used the Friedman test for the “Strangeness” and “Life-meaning” ratings, and a one-way repeated measures ANOVA for the remaining ratings, as determined by Shapiro-Wilk tests of normality. No significant main effects were observed for any of the items (Supplementary Figure 1-13).

### 3.5 Decoded urge to move

Bayesian GLMM results (Table 3), together with subsequent pairwise comparisons (Table 4), showed no differences in decoded “urge to move” between any playlists. All 95% HPD intervals for the contrasts included zero.

**Table 3:**
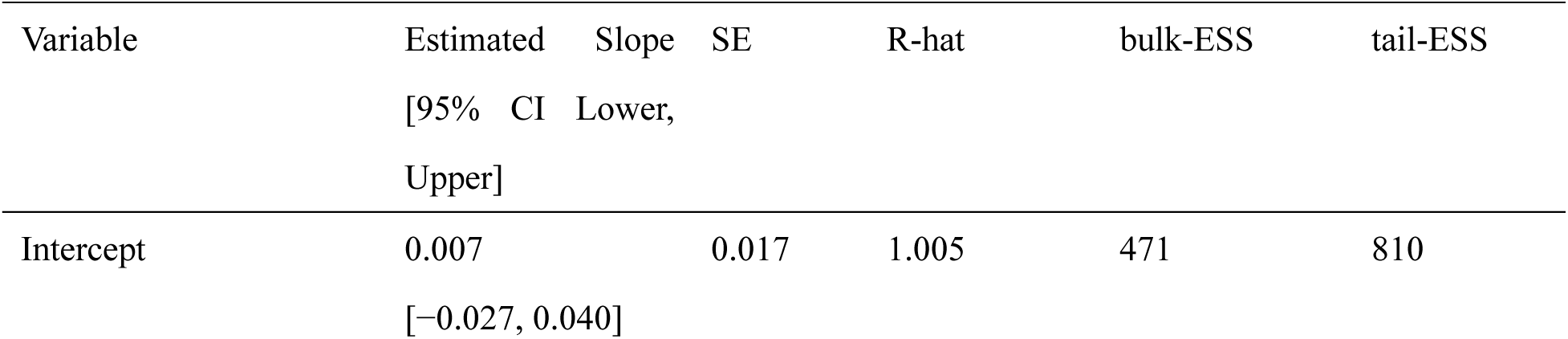
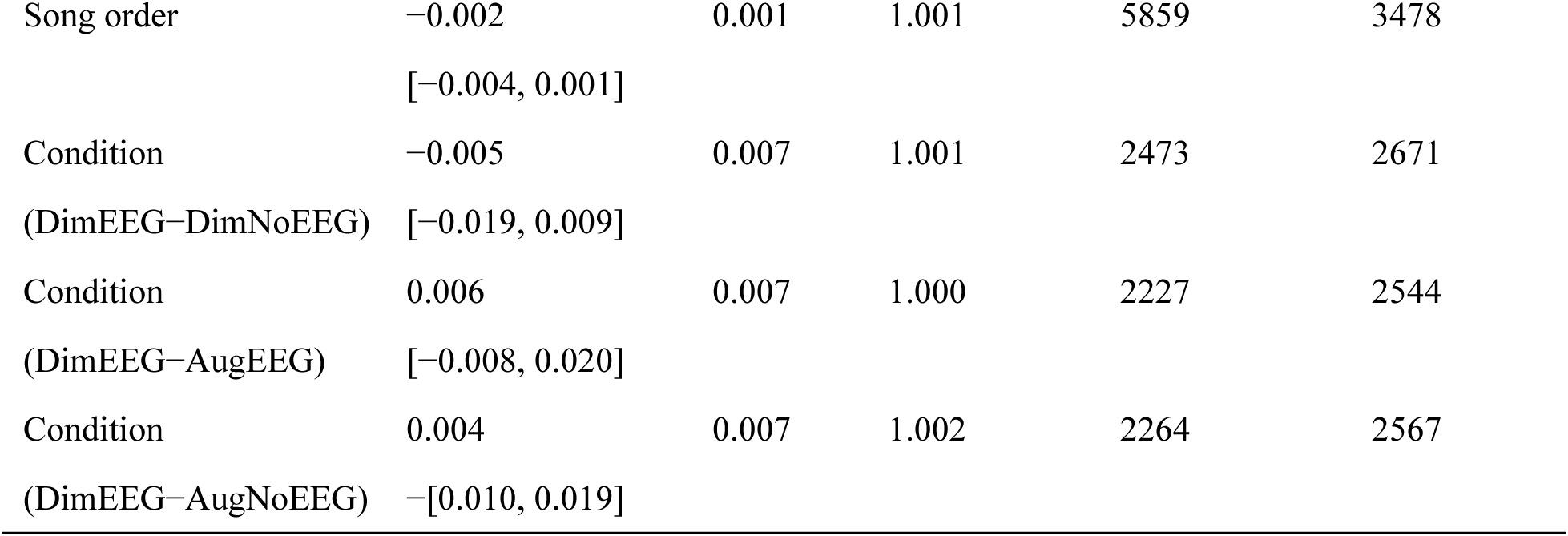
The results of Bayesian generalized mixed-effects model on the decoded urge to move.

**Table 4:** Results of pairwise contrasts of decoded “urge to move” across playlists, with 95% HPD intervals.

| Contrast | Estimate Difference [95% HPD Intervals:<br>Lower, Upper] |
| --- | --- |
| AugEEG–DimEEG | 0.006 [−0.008, 0.019] |
| AugEEG–DimNoEEG | 0.011 [−0.003, 0.025] |
| AugEEG–AugNoEEG | 0.002 [−0.012, 0.016] |
| AugNoEEG–DimEEG | 0.004 [−0.010, 0.018] |
| AugNoEEG–INVnoEEG | 0.009 [−0.004, 0.023] |
| DimEEG–DimNoEEG | 0.005 [−0.009, 0.019] |

### 3.6 Individual differences (BMRQ and Gold-MSI)

The stepwise procedure selected a model with three BMRQ factors: Sensory Motor, Music Seeking, and Mood Regulation (*F*(3, 18) = 3.87, adjusted *R*² = 0.29, *p* = 0.027; AIC = 130.46). In this model, Sensory Motor was a significant predictor of MoveDiffEEG (Table 5).

**Table 5:**
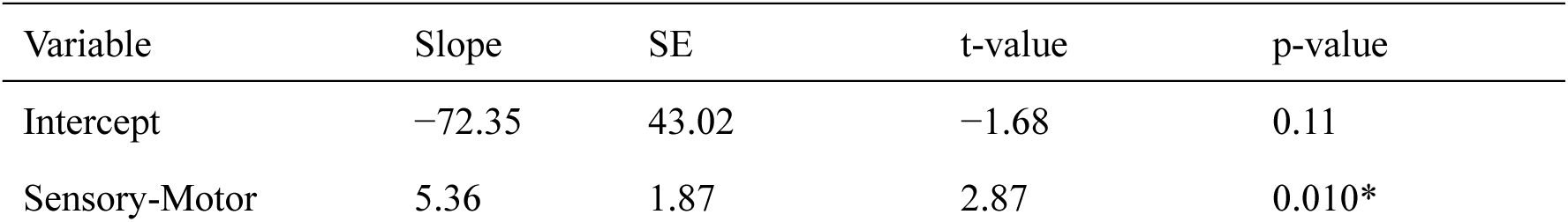
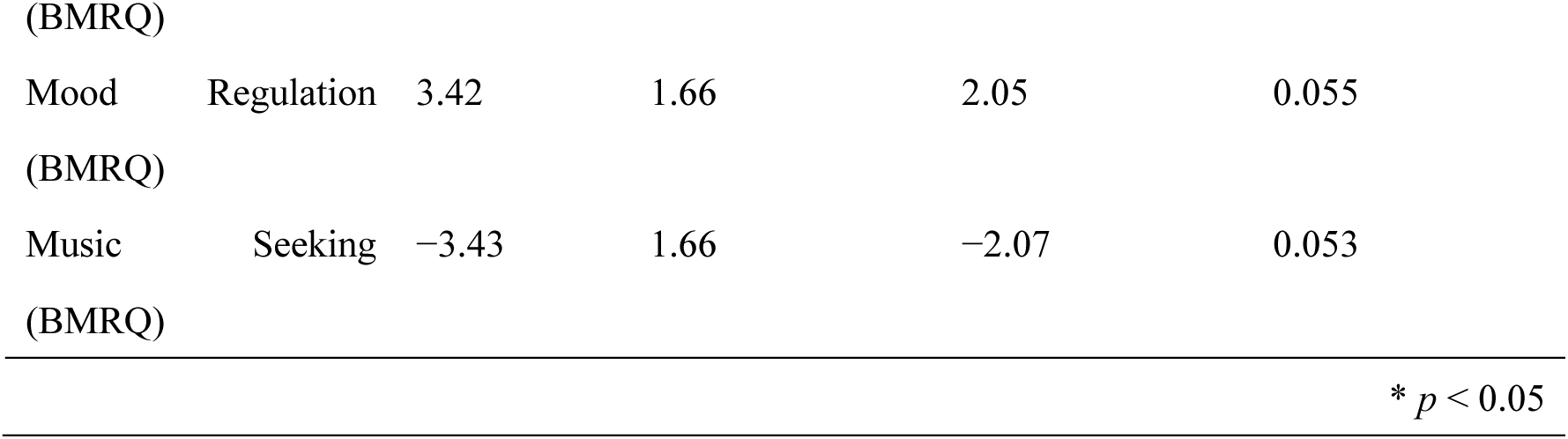
The results of stepwise regression analysis on MovDiffEEG. NovDiffEEG is defined as the difference in “urge to move” ratings between AugEEG and DimEEG (AugEEG minus DimEEG).

## 4. Discussion

### 4.1 Subjective ratings for the “urge to move” and “pleasure”

In this study, we aimed to develop a groove-brain-music interface (G-BMI) that uses in-ear EEG to generate personalized playlists that enhance each individual’s groove experience. AugEEG received higher “urge to move” ratings than DimNoEEG and DimEEG, and higher “groove” ratings (urge to move + pleasure) than DimEEG. These findings indicate that incorporating individual EEG data in real-time improves playlist generation compared with models that rely only on acoustic features.

Our results indicate that although music that induces groove shares several common musical characteristics, such as a moderate degree of syncopation, an optimal tempo, and strong beat salience as shown in previous studies (Etani et al., 2018; Madison et al., 2011; Senn et al., 2018; Stupacher et al., 2016; Witek et al., 2014), there are also individual differences across listeners. Furthermore, these results suggest that it is possible to build a high quality and high precision music recommendation system by incorporating biological data, including EEG. Wearable devices such as the in-ear EEG used in this study will be useful for implementing such a service.

There were no significant differences among the playlists in “pleasure” ratings. Although prior studies have shown a strong correlation between “urge to move” and “pleasure” (Etani et al., 2018; Witek et al., 2014), this result suggests that the two are indeed not identical. It is possible that constructing a model to induce pleasure is intrinsically more difficult. Nevertheless, a previous study succeeded in building such a model using a similar approach (Kondoh et al., 2024). Thus, the more plausible interpretation is that “urge to move” and “pleasure” are correlated but distinct constructs.

The total groove rating, defined as the sum of “urge to move” and “pleasure,” was also higher for AugEEG than for DimEEG. Thus, a model trained on “urge to move” ratings alone increased the overall groove rating. In this study, we used only the “urge to move” ratings because the common definition of groove as “the pleasurable urge to move to music” combines two distinct components, which makes direct modeling more difficult. It is possible that a G-BMI trained on the composite construct would enhance the overall groove experience more effectively than the present model. This should be examined in future studies.

### 4.2 Subjective well-being and emotion-related ratings

In this study, participants also rated each playlist on well-being and emotion related items. No significant differences were observed among playlists for any of these items. Research on the link between groove and well-being is limited. A previous study found a positive correlation between groove ratings and happiness ratings, suggesting that experiencing groove may increase happiness and contribute to mental well-being (Kawase & Eguchi, 2010). The same study also reported a positive correlation with a sense of unity. Although this does not directly assess well-being, it implies that experiencing groove may foster feelings of closeness to others and, in turn, social bonding that supports mental health. In contrast to these findings, in our data, enhancing groove experience or the urge to move did not by itself increase self-reported well-being. Because physical exercise benefits mental health (Pearce et al., 2022), we expected that listening to music that induces an urge to move might also improve well-being, but this was not supported. A plausible explanation is that experiencing groove alone is insufficient and actual movement in response to the urge to move may be necessary. In our protocol, participants were asked to sit still to obtain high quality EEG recordings, which may have attenuated any potential benefits. Allowing movement during listening could yield different outcomes. In addition, Kondoh et al. (2024) reported stress reduction when playlists were optimized to enhance pleasure. Thus, generating playlists with a model that targets the total groove experience, rather than the urge to move alone, may also facilitate stress reduction.

Regarding emotion-related items, previous studies have reported significant correlations between groove ratings and emotions such as cheerfulness (Kawase & Eguchi, 2010). However, playlists optimized for the “urge to move” did not affect emotion related ratings. The correlation coefficient for cheerfulness was 0.28, suggesting small to moderate associations that are not strong enough to induce emotions. In Kondoh et al. (2024), items such as absorption, excitement, arousal, and valence received higher ratings for pleasure-inducing playlists. Thus, targeting the composite groove experience, rather than the urge to move alone, may influence these emotions.

### 4.3 Decoded urge to move

As noted above, although the subjective “urge to move” and groove ratings differed across playlists, the decoded urge to move did not differ between playlists. This suggests that, while we successfully generated groove inducing playlists by updating the model and the ranking using in-ear EEG, the subjective differences were not fully captured by the decoded EEG measure. One plausible explanation is that, although the decoder was trained on EEG recorded while participants listened to excerpts with high or low “urge to move” ratings, those excerpts were originally selected as high-groove or low-groove stimuli from a previous study (Janata et al., 2012) and thus carry information beyond “urge to move,” including pleasure. As a result, the decoded “urge to move” score likely reflected broader properties of those stimuli rather than the pure “urge to move” construct. Even so, this decoded index was useful for guiding music selection, as evidenced by the subjective playlist ratings.

### 4.4 Individual differences (BMRQ and Gold-MSI)

Stepwise multiple regression, with MovDiffEEG (AugEEG −DimEEG) as the dependent variable and BMRQ and Gold MSI scores as independent variables, showed that the Sensory-Motor subscale score of BMRQ was significantly associated with MovDiffEEG. The positive coefficient indicates that participants with higher Sensory-Motor scores exhibited larger MovDiffEEG, consistent with greater sensitivity to the urge to move. Items on the Sensory-Motor factor include statements such as “Music often makes me dance,” so this association is plausible. In contrast, no Gold-MSI subscales showed significant effects. This pattern suggests that sensitivity to the urge to move while listening to music depends less on general musical sophistication and more on movement related responsiveness captured by the BMRQ.

### 4.5 Limitations

Our study has several limitations. First, although we succeeded in enhancing groove experience using individual EEG data, the decoded EEG did not differ across playlists. Therefore, it is difficult to conclude that differences in subjective ratings were reflected in the decoded EEG measures. Nevertheless, our results suggest that updating the model and the ranking based on EEG was useful for recommending high-groove music on an individual basis. Second, as noted above, we used only the “urge to move” ratings when developing the model. Consequently, although the total groove rating was higher for AugEEG than for DimEEG, the difference was smaller than for “urge to move” alone.

To better maximize the overall groove experience, future work should train the model directly on the total groove ratings. Third, although we aimed to develop an individualized G-BMI, participants in the present study were relatively young on average (mean age = 24.55 ± 5.91 years). To confirm the system’s validity and generalizability, future work should evaluate its accuracy in samples spanning a broader age range.

## 5. Conclusion

In this study, we aimed to develop a closed-loop G-BMI that uses individual EEG data to maximize the groove experience. The playlist optimized to increase groove, selected using both acoustic features of musical excerpts and real-time in-ear EEG, received higher “urge to move” and total “groove (urge to move + pleasure)“ ratings than the playlist optimized to decrease groove using the same information. These findings suggest that our closed-loop neurofeedback system leveraging individual EEG successfully recommended high-groove music tailored to each participant, and that such systems can help maximize individual musical experiences.

**Supplementary Material**

## Methods

### 1. Recording

Participants sat in front of the PC, and wore in-ear EEG device (VIE Inc., Japan), and tuned the volume. They listened to six musical excerpts (three high-groove, and three low-groove musical excerpts selected from Janata et al. (2012), 90 seconds each) and rated the “urge to move” and “pleasure” using a VAS. While they listened to musical excerpts, we recorded participants’ in-ear EEG. EEG data were recorded from the canals, and the reference electrode was placed on the back of the neck. Electrically conducted ear tips were attached to the electrodes inserted into the ear canals, which also functioned as earphones. The sampling rate was set to 600 Hz. While listening to the musical excerpts, participants were instructed to stay relaxed, avoid from moving their body, and gaze at a cross attached on a wall in front of them (about 1 m away). After the presentation of each musical excerpt, they rated the “urge to move” and “pleasure” using a VAS (0-100). This section lasted about 20 minutes.

## 2. Modeling

### 2.1 Model 1

We trained a LASSO regression model to predict “urge to move” ratings from acoustic features of the musical excerpts. Acoustic features were computed with VGGish, a pretrained neural network. For each participant, VGGish converted six excerpts (three high-groove and three low-groove, 90 seconds each) into 372 time frames, each represented by a 128 dimensional embedding. We then averaged the features within nonoverlapping 10 second windows, yielding 9 time bins per excerpt. This produced a feature matrix of 6 excerpts by 9 time bins by 128 features.

We computed the mean and standard deviation (SD) of each feature across the six musical excerpts for each dimension, and standardized the acoustic features to z-scores. To improve model accuracy, we retained only feature dimensions without missing values and with SD greater than 0.01. Finally, we fit a linear LASSO model with the z-scored “urge to move” ratings as the response variable, and the retained z-scored acoustic features as predictors, as described by the following formula.

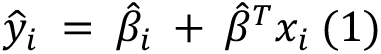

Where 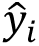 is the predicted subjective “urge to move” rating for musical excerpt 𝑖. 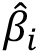 is the intercept, and 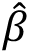 is the vector of coefficients, one for each acoustic feature dimension. 𝑥_i_ is the vector of selected acoustic feature values calculated by VGGish for musical excerpt 𝑖, with one value for each dimension. The parameters 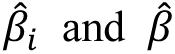 are the values of 𝛽_0_ and 𝛽, respectively, that minimize the following loss function:

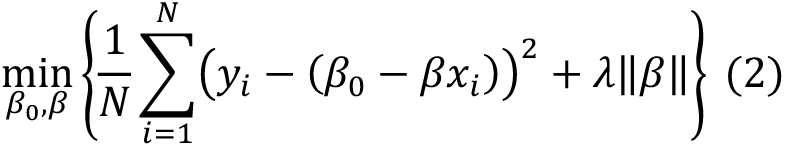

Where 𝜆 is the regularization parameter. The optimal 𝜆 was selected using fivefold cross-validation.

𝑁 denotes six, the number of musical excerpts used during the recording.

### 2.2 Model 2

To develop a model that classifies EEG signals into high or low “urge to move” states, we first tested whether participants reported greater “urge to move” while listening to high groove music than low groove music (t test; see Results).

For EEG preprocessing, the raw data were band pass filtered with a fourth order Butterworth filter from 3 to 40 Hz. We then performed independent component analysis (ICA) on the filtered signals from both electrodes and obtained two components. We computed the root mean square (RMS) of each component, labeled the component with the larger RMS as noise, and removed it. The data were segmented into four second windows with 50 percent overlap. For each epoch, we calculated the power spectral density from 4 to 40 Hz in 0.5 Hz steps. We also calculated the total power across all frequencies and the RMS of the filtered signal for each channel. In addition, we computed the maximum absolute value of the numerical gradient of the signal and the skewness. Each noise metric was standardized and then averaged across the right and left electrodes for each window, and a noise flag was assigned to windows that exceeded a predetermined threshold of 2.5. Only epochs without noise flags were used.

The EEG data were standardized across all epochs, frequency windows, and electrodes (left, right, and their difference). Principal component analysis (PCA) was then performed on the standardized EEG data. We retained at most 150 principal components; if fewer than 150 components were available, all available components were used. The mean and SD of the principal component scores were calculated and standardized. Each epoch was labeled according to whether it was recorded while listening to high-groove music (true) or low-groove music (false). When the numbers of true and false epochs were unequal, we randomly sampled from the larger group to match the smaller group. The data were split into training and test sets, with ninety percent used for training and ten percent for testing.

We then fit a logistic LASSO regression model to decode “urge to move” from EEG frequency components. The model’s predicted odds of high groove music, denoted 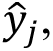 is expressed as follows.

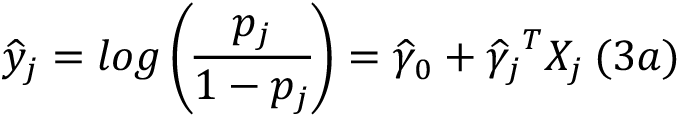

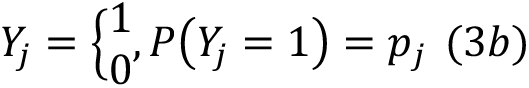

Where 𝑝_j_ is the probability that the EEG data are classified as being recorded while listening to high-groove music, 𝑗 is the number of EEG components, 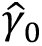 is the intercept, and 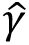 is the coefficient vector, with one coefficient for each EEG component. 𝑋_*_ denotes the vector of EEG component values. Note that 𝑌_*_ is a binary outcome variable. The parameters 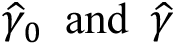 are the values of 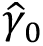 and 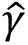 that minimize the following loss function:

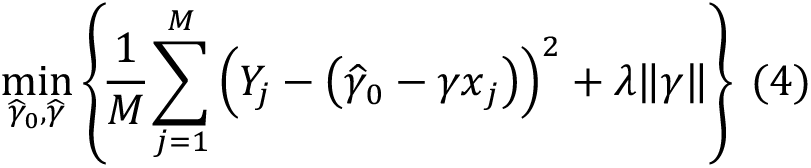

where 𝜆 is the regularization parameter. The optimal 𝜆 was selected using twenty-fold cross validation. 𝑀 denotes the maximum value among the EEG components.

We then computed the confusion matrix (true positives, false positives, false negatives, and true negatives) and the accuracy for the training set, as well as the receiver operating characteristic (ROC) curve and the area under the curve (AUC). The model was trained on the training set and evaluated on a separate test set, for which we calculated the same metrics, including the ROC curve and AUC. The mean accuracy rates for the training and test sets were 81.1% and 68.6%, respectively (Figure S14), and the mean AUC for the trained and test data were 87.3% and 72.6%, respectively (Figure S15). As shown earlier, participants experienced a stronger “urge to move” when listening to high-groove music.

Therefore, this generalized logistic LASSO classifier that distinguishes true and false epochs can be used to estimate the “urge to move” from EEG.

## 3. Generating playlists

### 3.1 Songs

The candidate pool comprised 7,225 songs drawn from the GfK Japan weekly music chart Top 1-1000 between April 2018 and February 2022. Acoustic features for each song were extracted with VGGish. Feature values were standardized across songs, and values exceeding two SD were clipped. We then assembled a feature matrix for Model 1 to predict subjective “urge to move.” For each candidate, we computed correlation coefficients between its feature values and those of high-groove music, converted these coefficients to z scores, and summed them to obtain a score. Candidates were ranked in descending order based on this score.

### 3.2 In-ear EEG measurement and decoding

EEG was recorded during playlist listening at a sampling rate of 600 Hz and a frequency range of 4-40 Hz. Signals were band pass filtered from 3 to 40 Hz using a Butterworth filter. From the filtered EEG data, we computed power spectra, normalized them, and entered them into Model 2 to predict “urge to move.” For noise detection, we calculated RMS, the maximum numerical gradient, and kurtosis, and assigned noise flags using the same criteria as in the previous modeling section. We then computed a moving average of the decoded “urge to move” state using a five second window with updates every second. When fewer than five valid points were available, we averaged all unflagged points. Otherwise, we averaged the most recent five unflagged points.

### 3.3 Procedure

Each participant listened to four playlists, each containing seven songs. Each song was 60 seconds long. The first song, used as the baseline, was randomly selected from ranks 3,577 to 3,649. At the end of each song, 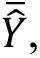 the 60 second average of the Model 2 predicted “urge to move” excluding noise segments, was converted to an approximate VAS value, 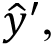 as follows.

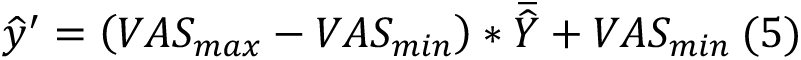

where 𝑉𝐴𝑆_max_ and 𝑉𝐴𝑆_min_ are the maximum and minimum VAS ratings in the first recording section, respectively. 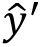 was standardized using the mean and SD of the VAS ratings. The acoustic features of the song was precalculated using VGGish and converted to z-scores. We then added the standardized decoded “urge to move” level from the EEG and acoustic features of the played songs to Model 1 and retrained the model. This procedure was repeated until the playlist was complete.

The four playlists comprised two that augmented the “urge to move” (AugEEG and AugNoEEG) and two that diminished it (DimEEG and DimNoEEG). For AugEEG, the updated Model 1 produced an “urge to move” prediction score for each candidate song. This score was combined with the z-score of the correlation between the song’s feature values and those of high-groove music, and candidates were then re-ranked in descending order to update the ranking. This playlist drew from the top 72 songs (top 1% of all songs). The other augmenting playlist, AugNoEEG, was not updated with the retrained Model 1 and remained fixed based on the initial ranking without EEG. To prevent overlap between the two augmenting playlists, AugNoEEG drew from the top 432 songs (top 6%). As AugEEG repeatedly selected from the top 72 in six iterations, it yielded 432 selections in total (allowing duplicates), which matches the candidate pool size for AugNoEEG.

The two diminishing playlists followed the same logic. DimEEG, the EEG updated playlist, drew from the bottom 72 songs (bottom 1%), whereas DimNoEEG drew from the bottom 432 songs (bottom 6%). Like AugEEG, DimEEG repeated selection from the bottom 72 across six iterations.

At the end of each playlist, participants rated the entire playlist on 14 items (see Table 2), including “urge to move” and “pleasure,” using a VAS from 0 to 100. After listening to all playlists, they completed the Japanese versions of the Barcelona Music Reward Questionnaire (BMRQ) and the Goldsmiths Musical Sophistication Index (Gold-MSI).

## Supplementary Figures

**Figure S1:**
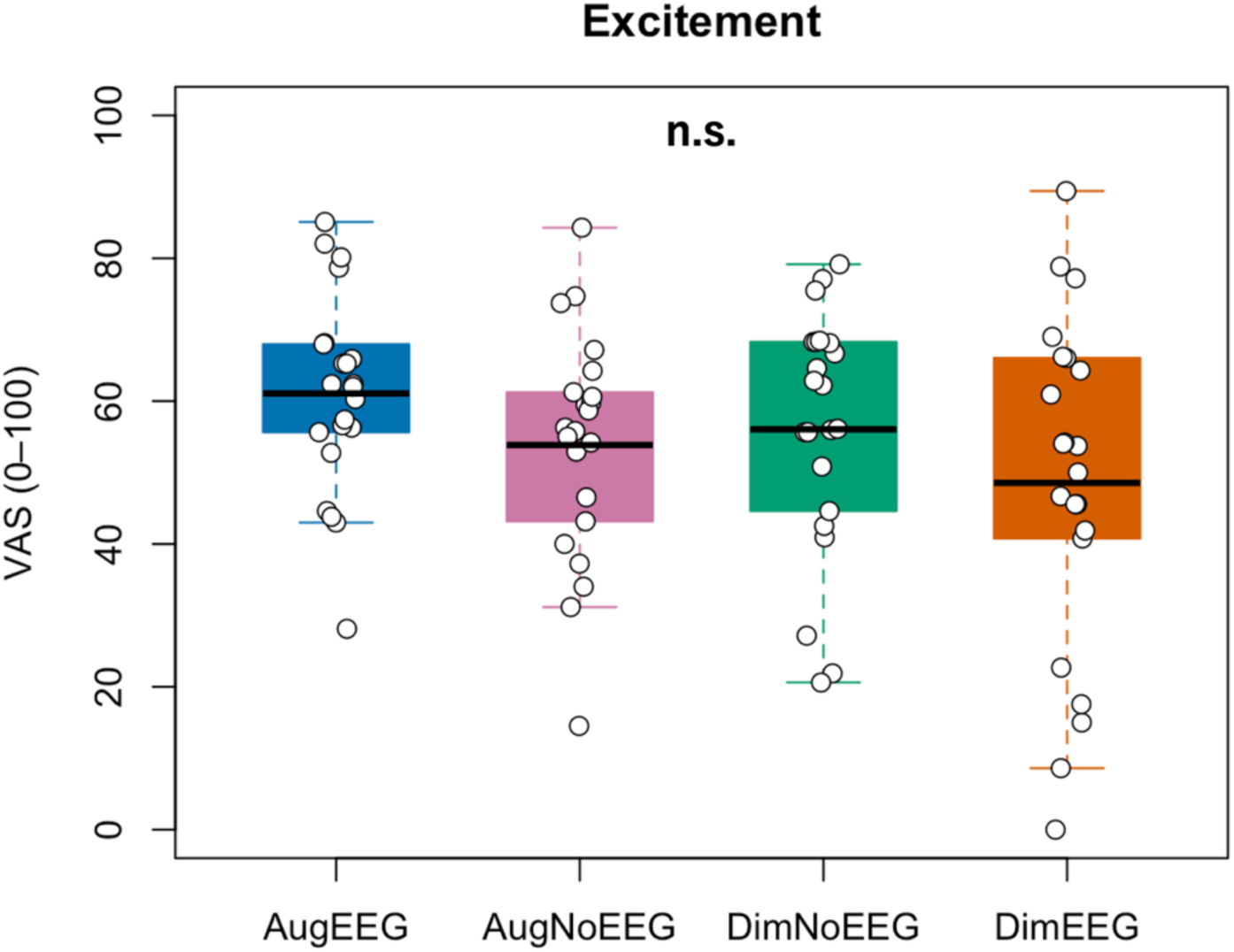
Subjective “Excitement” ratings for each playlist (see Table 2 for the full question).

**Figure S2:**
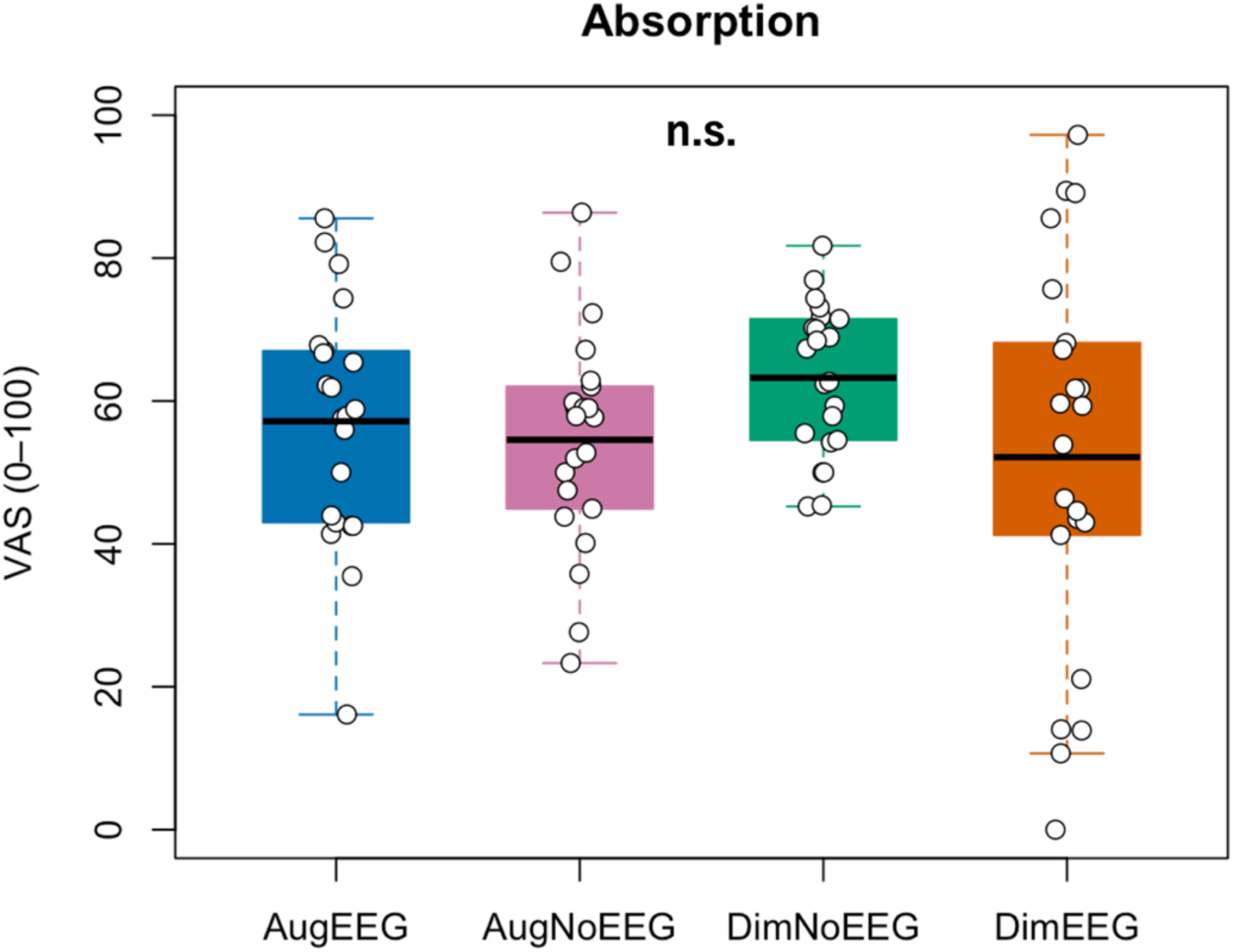
Subjective “Absorption” ratings for each playlist (see Table 2 for the full question).

**Figure S3:**
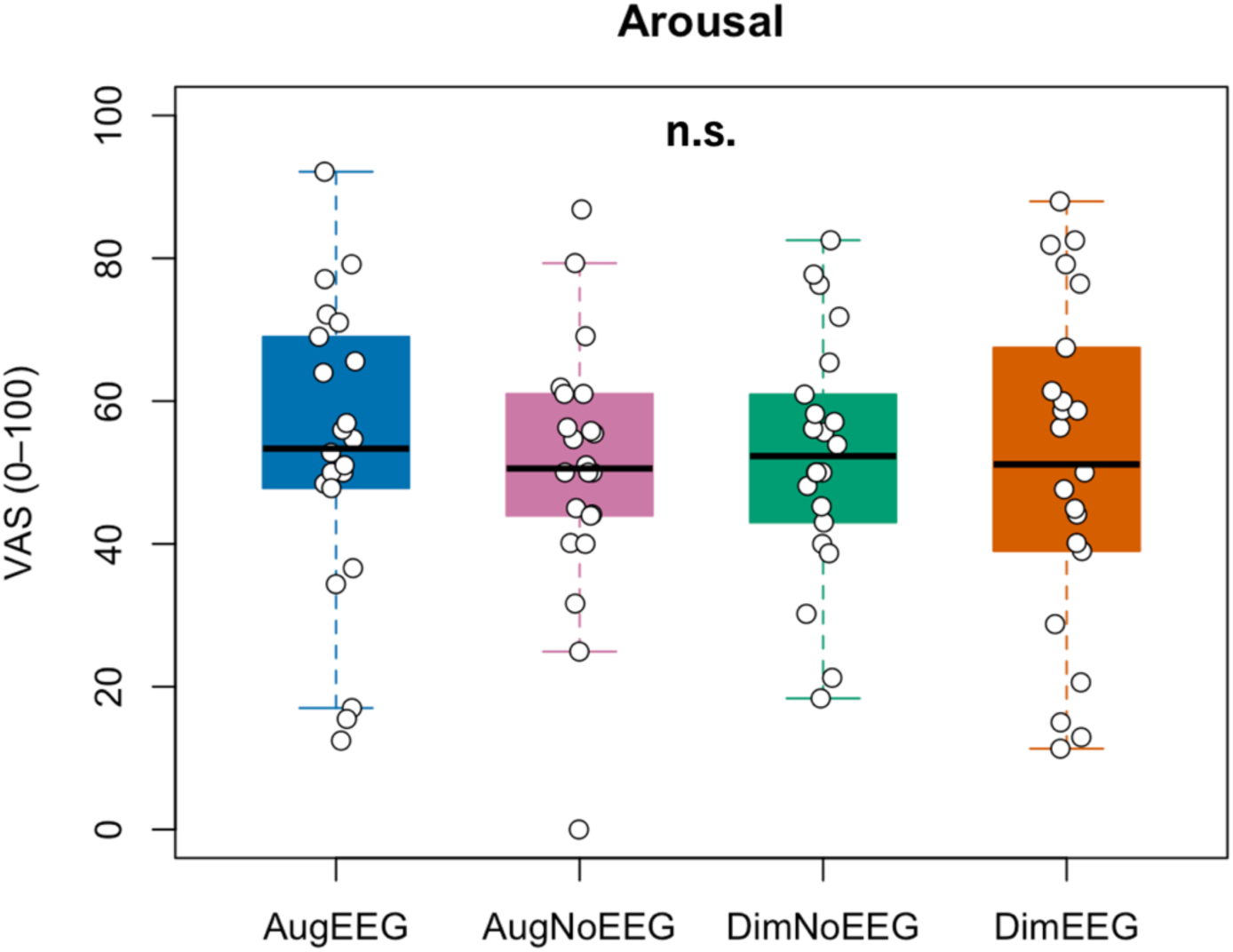
Subjective “Arousal” ratings for each playlist (see Table 2 for the full question).

**Figure S4:**
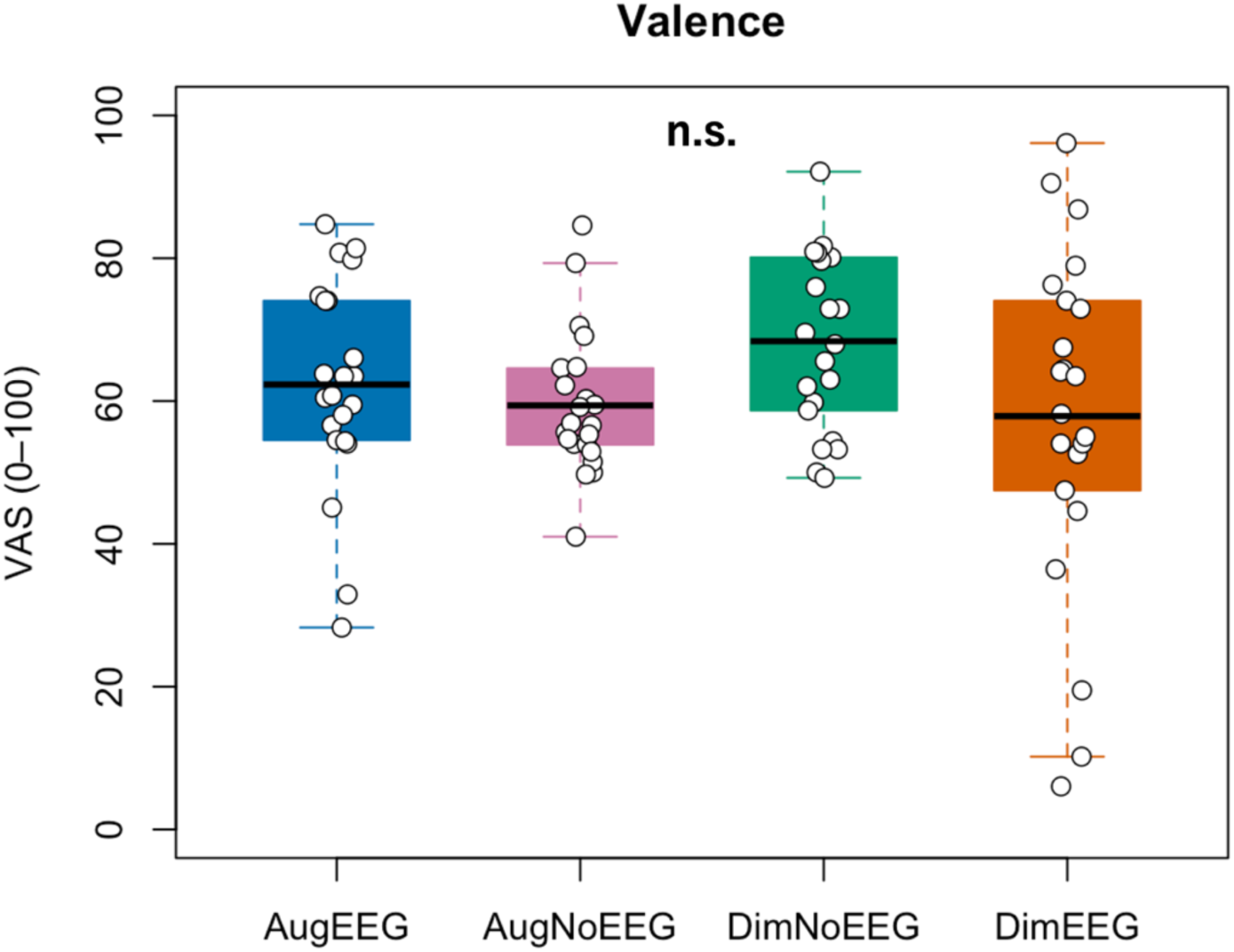
Subjective “Valence” ratings for each playlist (see Table 2 for the full question).

**Figure S5:**
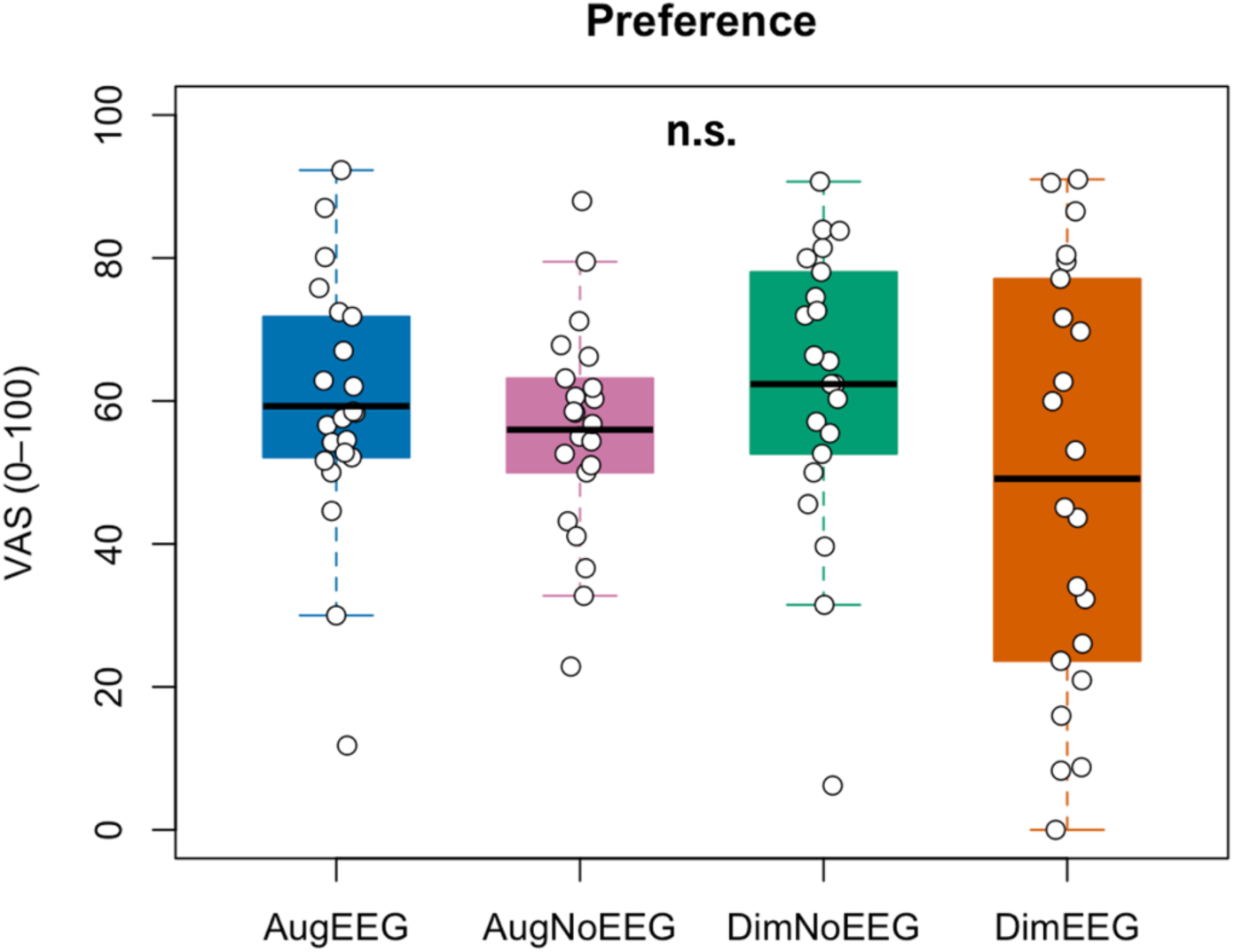
Subjective “Preference” ratings for each playlist (see Table 2 for the full question).

**Figure S6:**
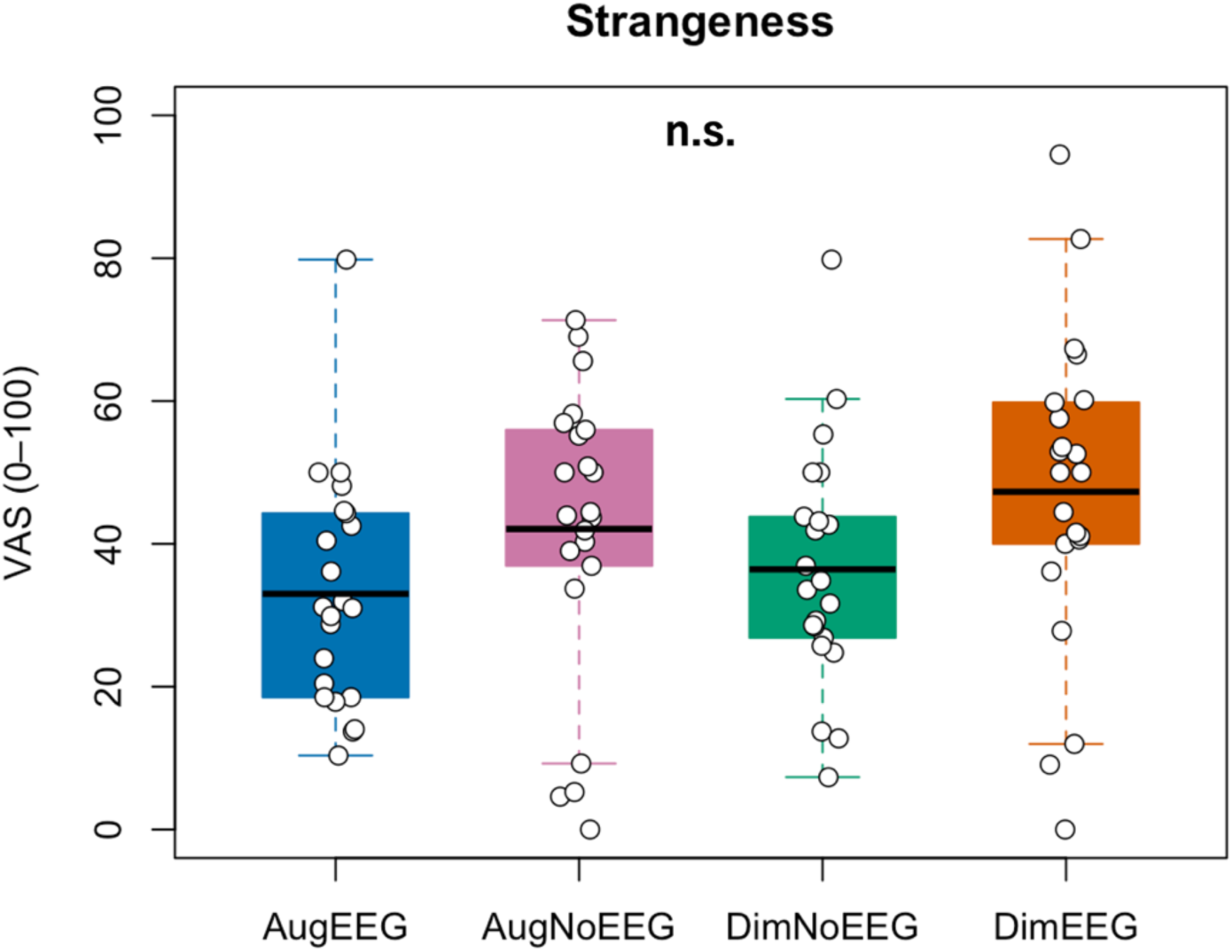
Subjective “Strangeness” ratings for each playlist (see Table 2 for the full question).

**Figure S7:**
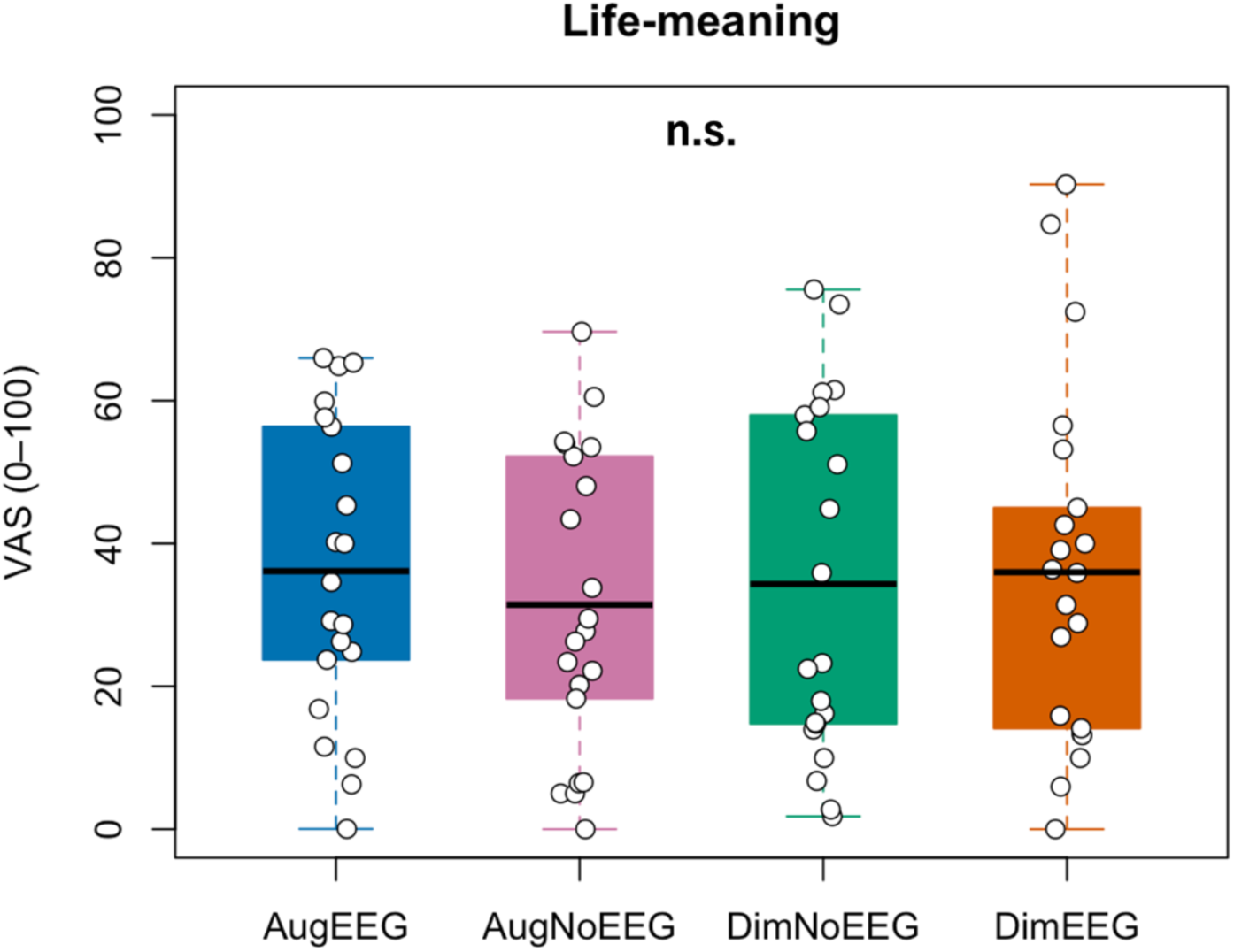
Subjective “Life-meaning” ratings for each playlist (see Table 2 for the full question).

**Figure S8:**
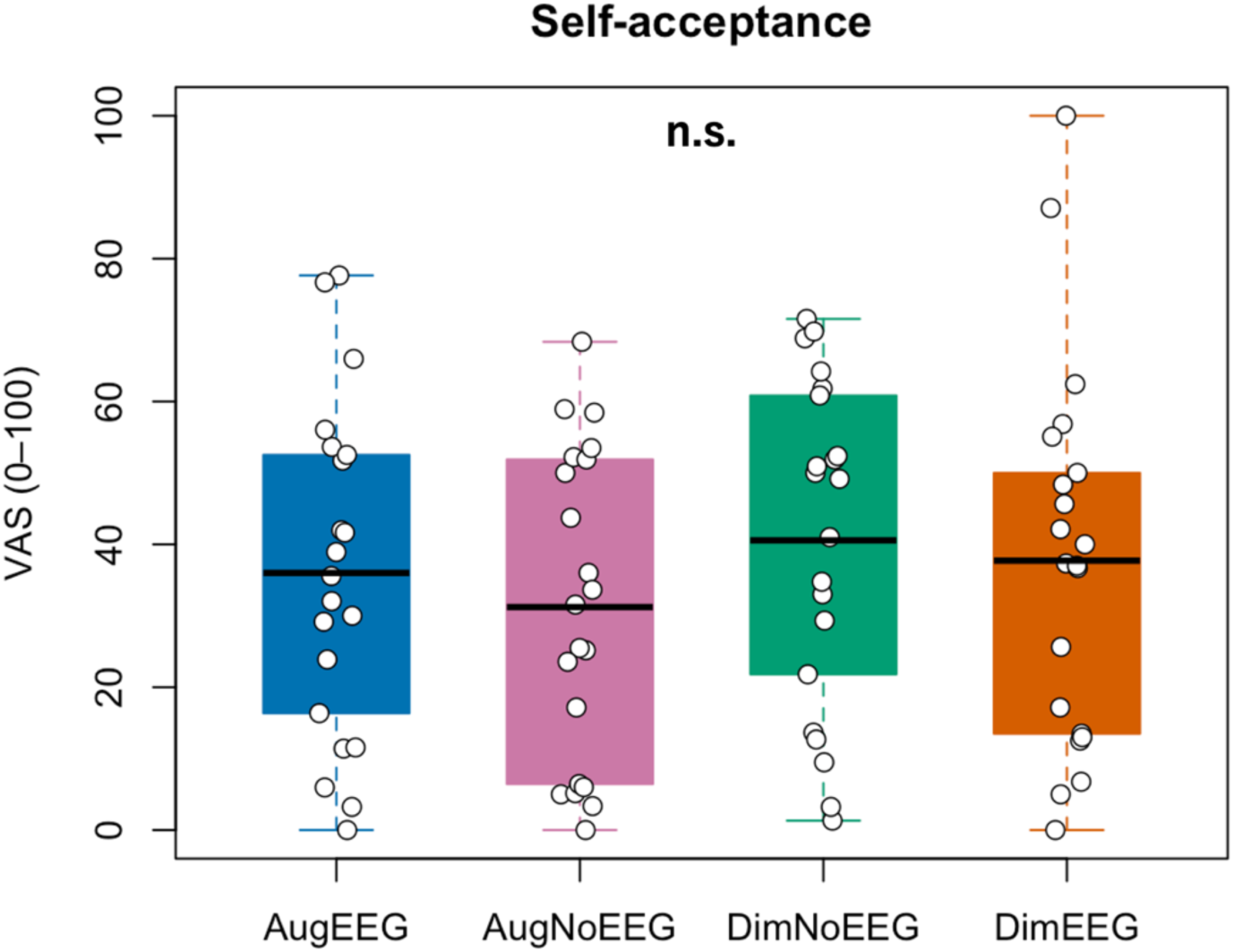
Subjective “Self-acceptance” ratings for each playlist (see Table 2 for the full question).

**Figure S9:**
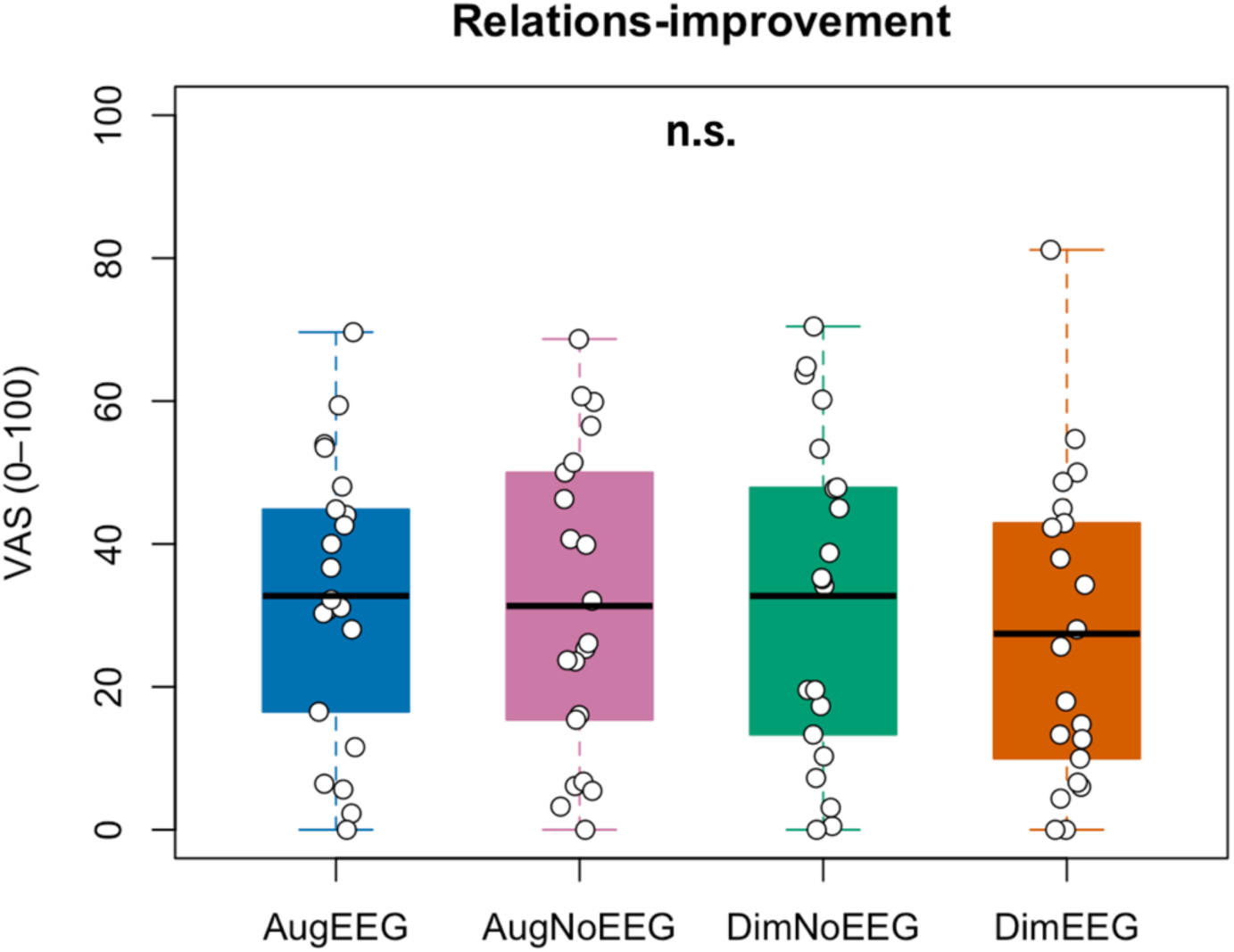
Subjective “Relations-improvement” ratings for each playlist (see Table 2 for the full question).

**Figure S10:**
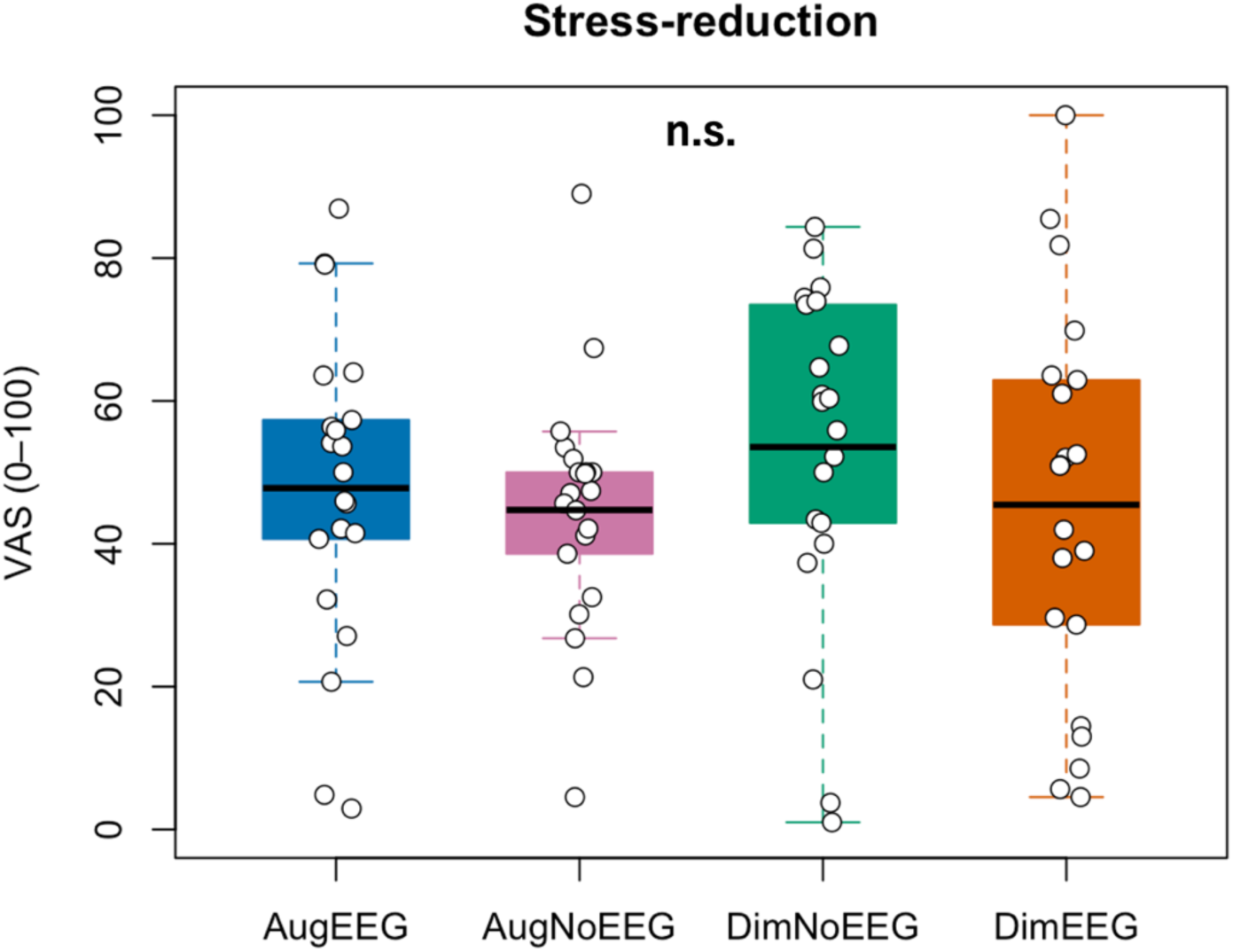
Subjective “Stress-reduction” ratings for each playlist (see Table 2 for the full question).

**Figure S11:**
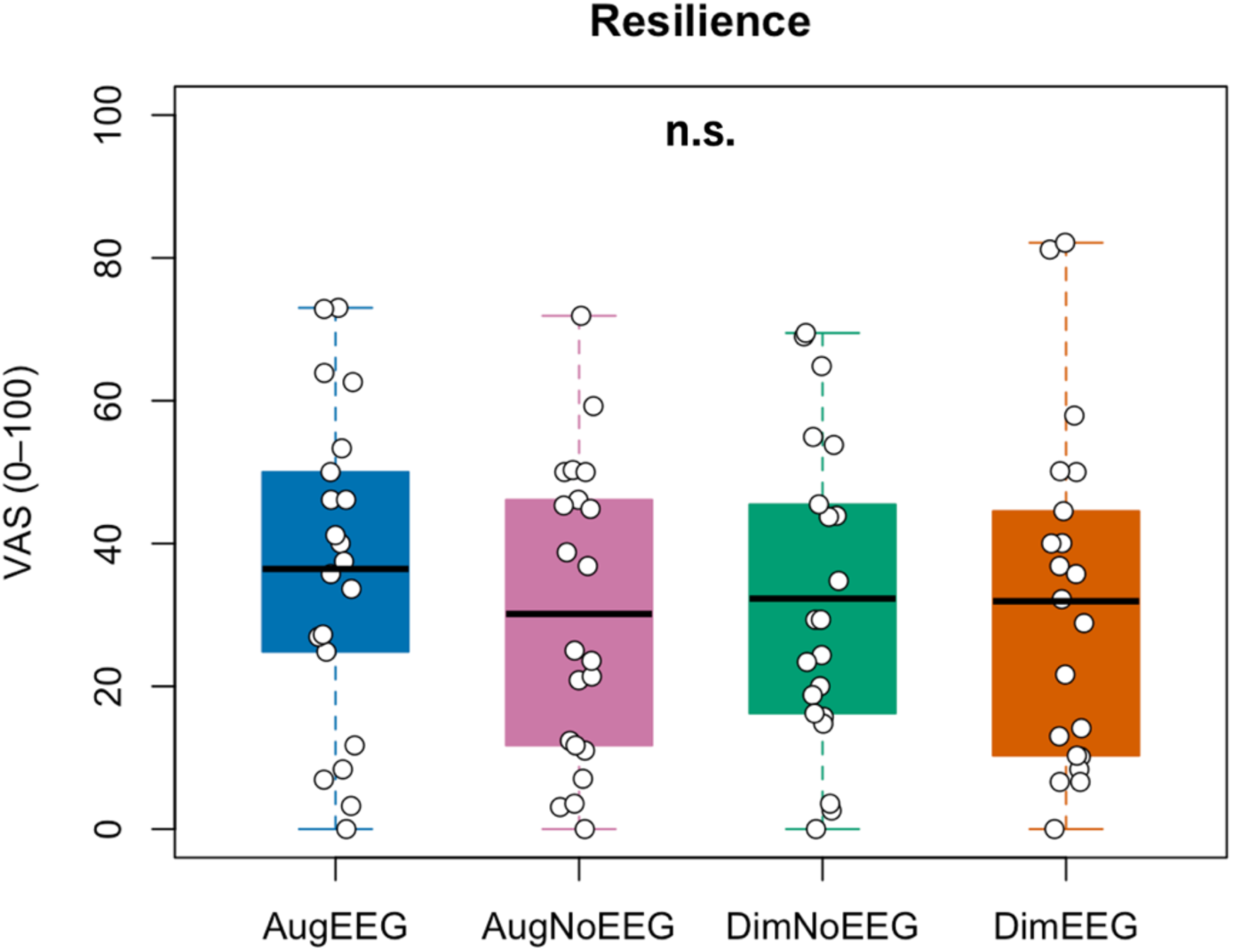
Subjective “Resilience” ratings for each playlist (see Table 2 for the full question).

**Figure S12:**
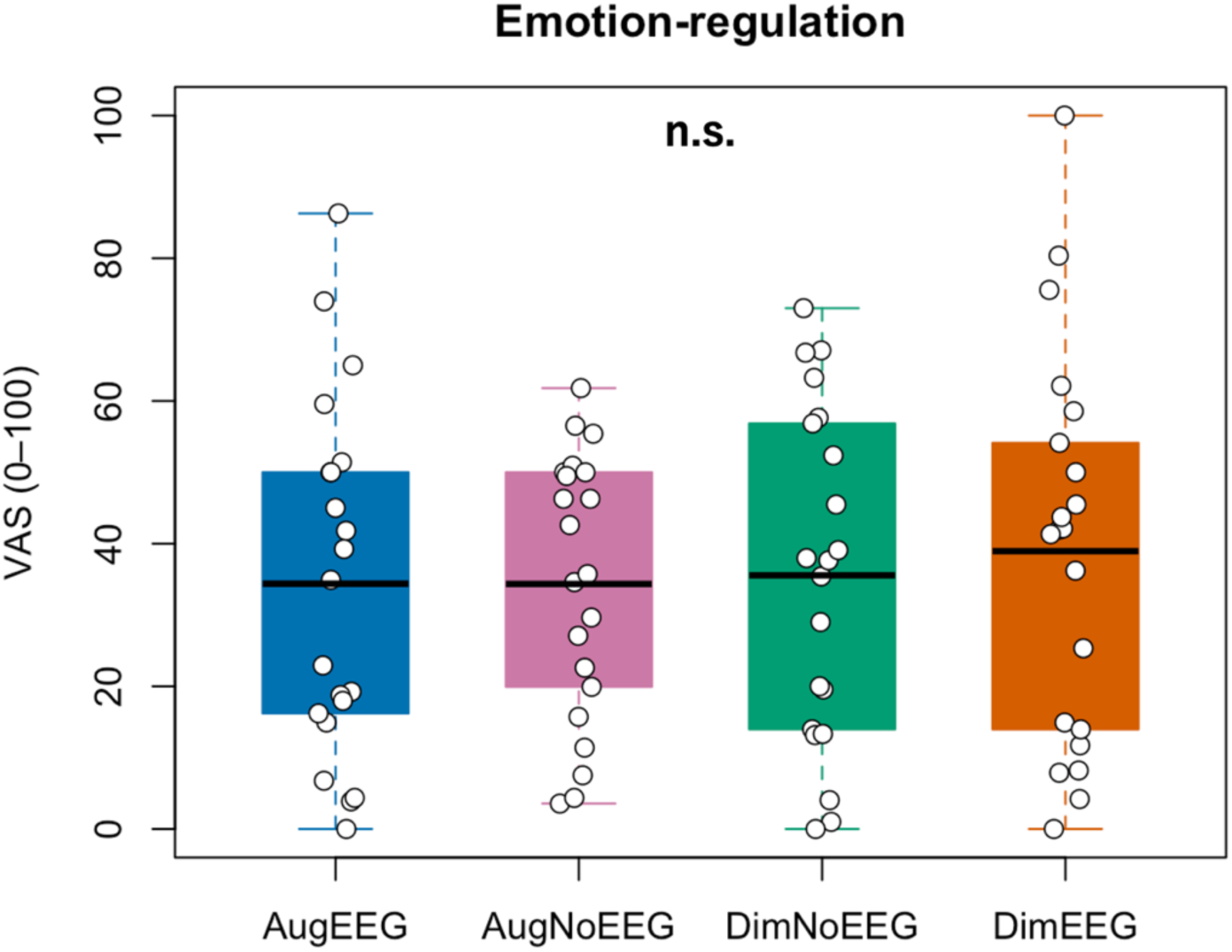
Subjective “Emotion-regulation” ratings for each playlist (see Table 2 for the full question).

**Figure S13:**
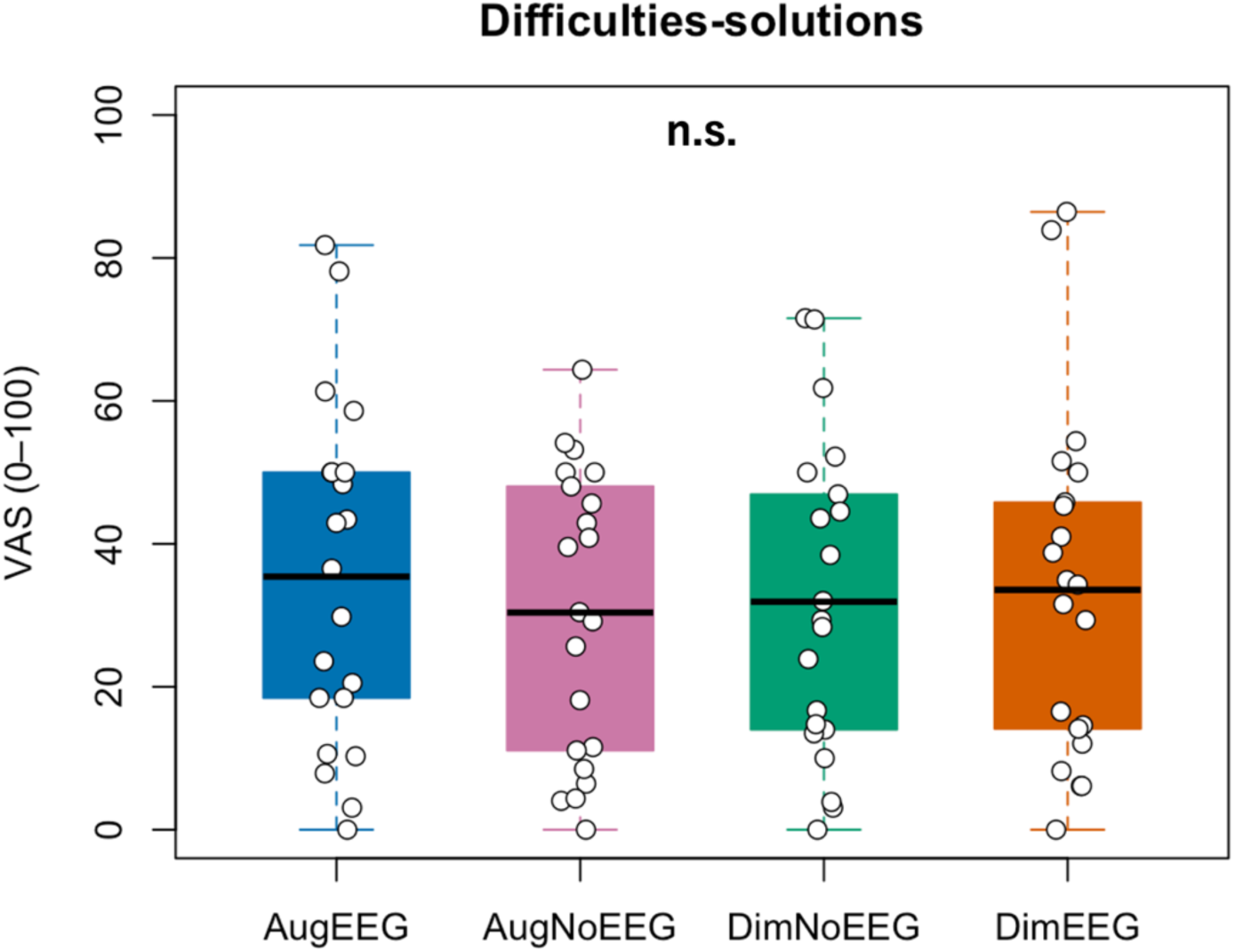
Subjective “Difficulties-solutions” ratings for each playlist (see Table 2 for the full question).

**Figure S14:**
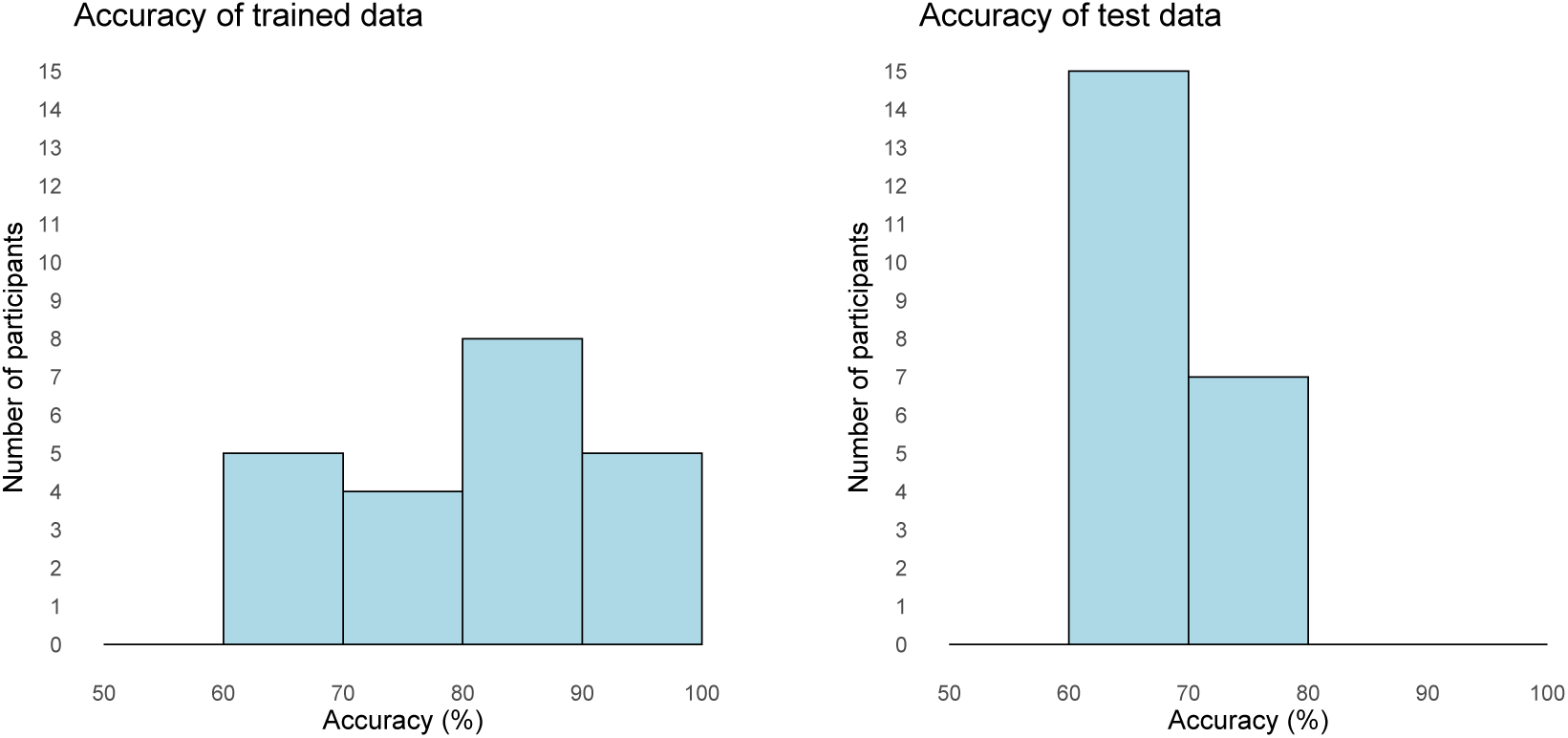
Histograms of the accuracy of trained and test data (*n* = 24).

**Figure S15:**
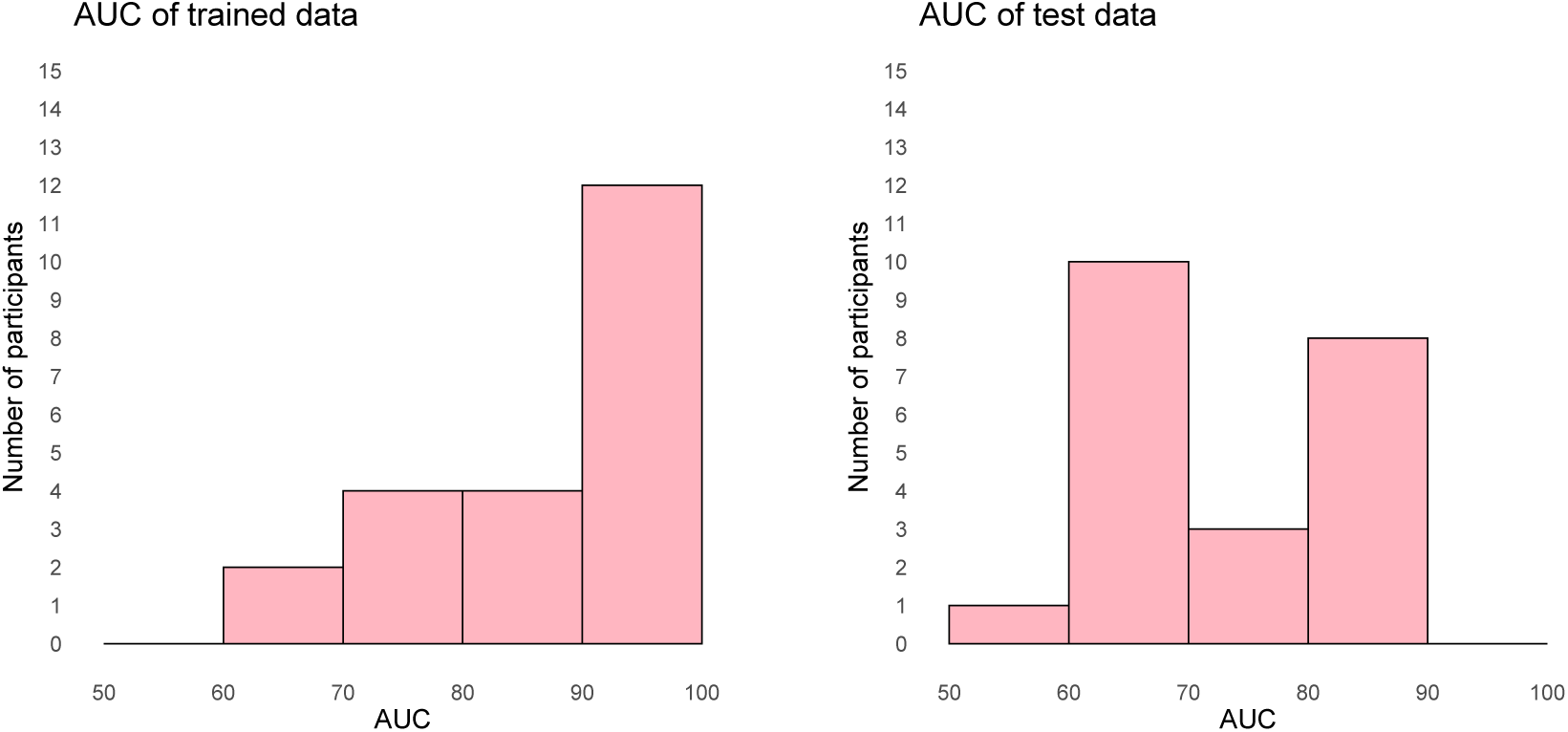
Histograms of the AUC of trained and test data (*n* = 24).

